# Bioequivalence Assessment of High-Capacity Polymeric Micelle Nanoformulation of Paclitaxel and Abraxane^®^ in Rodent and Non-Human Primate Models Using a Stable Isotope Tracer Assay

**DOI:** 10.1101/2021.08.20.457164

**Authors:** Duhyeong Hwang, Natasha Vinod, Sarah L. Skoczen, Jacob D. Ramsey, Kelsie S. Snapp, Stephanie A. Montgomery, Mengzhe Wang, Chaemin Lim, Jonathan E. Frank, Marina Sokolsky-Papkov, Zibo Li, Hong Yuan, Stephan T. Stern, Alexander V. Kabanov

**Author notes:** Authors contributed equally on this study (Duhyeong Hwang, Natasha Vinod). Corresponding Authors: Center for Nanotechnology in Drug Delivery, UNC Eshelman School of Pharmacy, University of North Carolina at Chapel Hill, 125 Mason Farm Road, Marsico Hall, Office #2012, Campus Box 7362, Chapel Hill, NC 27599-7362, USA.

## Abstract

The *in vivo* fate of nanoformulated drugs is governed by the physicochemical properties of the drug and the functionality of nanocarriers. Nanoformulations such as polymeric micelles, which physically encapsulate poorly soluble drugs, release their payload into the bloodstream during systemic circulation. This results in three distinct fractions of the drug-nanomedicine: encapsulated, protein-bound, and free drug. Having a thorough understanding of the pharmacokinetic (PK) profiles of each fraction is essential to elucidate mechanisms of nanomedicine-driven changes in drug exposure and PK/PD relationships pharmacodynamic activity. Here, we present a comprehensive preclinical assessment of the poly(2-oxazoline)-based polymeric micelle of paclitaxel (PTX) (POXOL *hl*-PM), including bioequivalence comparison to the clinically approved paclitaxel nanomedicine, Abraxane^®^. Physicochemical characterization and toxicity analysis of POXOL *hl*-PM was conducted using standardized protocols by the Nanotechnology Characterization Laboratory (NCL). The bioequivalence of POXOL *hl*-PM to Abraxane^®^ was evaluated in rats and rhesus macaques using the NCL’s established stable isotope tracer ultrafiltration assay (SITUA) to delineate the plasma PK of each PTX fraction. The SITUA study revealed that POXOL *hl*-PM and Abraxane^®^ had comparable PK profiles not only for total PTX but also for the distinct drug fractions, suggesting bioequivalence in given animal models. The comprehensive preclinical evaluation of POXOL *hl*-PM in this study showcases a series of widely-applicable standardized studies by NCL for assessing nanoformulations prior to clinical investigation.

**GRAPHICAL ABSTRACT:** 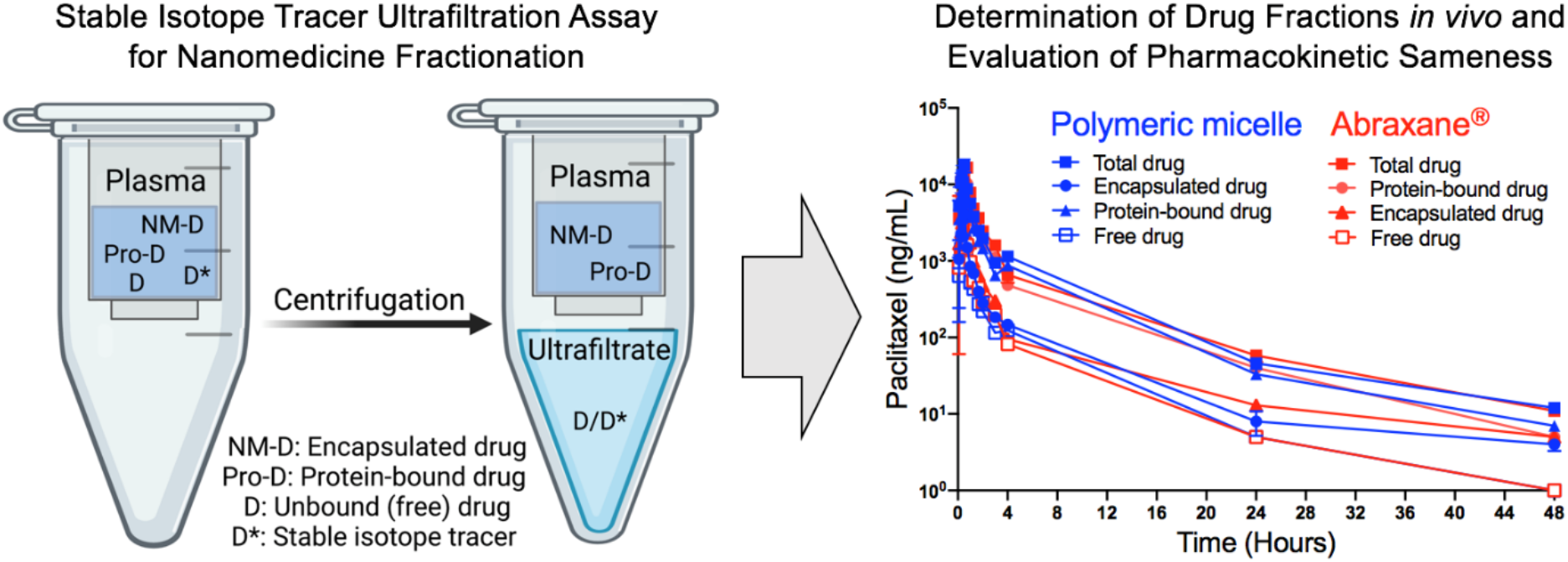

## Introduction

The pharmacokinetic (PK) evaluation of medicines provides valuable information about drug disposition and is key in predicting therapeutic and toxicity outcomes [1]. The PK evaluation of drugs formulated into nanocarriers (“nanoformulated drugs”) is challenging because their PK profiles are more complex due to the presence of multiple drug forms. While traditional drug formulations typically have two drug fractions, protein-bound and unbound (free) drug, with the advent of nanoformulated drugs, a third fraction that is nanoparticle encapsulated drug adds another layer of complexity to the PK analysis [2]. Furthermore, most PK studies are limited in their approach in that they depend on the total plasma drug concentration to deduce the PK parameters of the drug, when in reality, it is only the free drug fraction that is capable of pharmacodynamic activity throughout the body [3]. In the case of nanoformulated drugs, PK analysis of total drug is likely to be misleading when trying to predict therapeutic outcome. This is because, while drugs encapsulated in nanocarriers often make up the majority of the systemic total drug, this fraction is not considered therapeutically active. The encapsulated fraction is, however, susceptible to additional transport phenomena and capable of controlled-release of the drug, ultimately altering overall drug biodistribution [3]. Thus, reliance on the total drug measurement for PK analysis of nanoformulated drugs is inadequate for interpreting their dose-response relationship.

Polymeric micelles represent a unique class of nanoformulated drugs due to their dynamic structural nature. Physical encapsulation of poorly soluble drugs through non-covalent molecular interactions between amphiphilic block copolymers and drugs is a promising polymeric micelle formulation strategy [4]. While stably harboring poorly soluble drugs in the core of the micelle, polymeric micelles release the payload spontaneously either through diffusion or following the dissociation of micelles once the concentration of the polymer is below the critical micelle concentration (CMC) [5]. Upon release from the micelle, a significant fraction of the poorly soluble drug binds to plasma proteins, most notably albumin, and drug bound plasma proteins often function as an innate reservoir/carrier for the poorly soluble drug [6, 7]. The unbound drug, which is the portion of released drug not bound to plasma proteins, remains dynamic and exchanges with the protein-bound drug [8, 9]. The extent of drug encapsulation in nanocarriers and protein binding of the drug influences the exposure of pharmacodynamically active unbound drug (clinically relevant fraction) to target tissues. However, it is also important to consider that the drug could be transported in the polymeric micelle directly to the tissues and released from the micelle in the tissue. Thus, nanocarrier encapsulation of drugs has the potential to alter PK and distribution of the drug, as well as drug partitioning across biological barriers [10–13]. It is, therefore, important to study the PK profiles of individual drug subpopulations to achieve a meaningful PK assessment of nanoformulated drugs.

Notably, there is a major need for procedures that can accurately assess the PK of different drug fractions [14]. One such procedure, the stable isotope tracer ultrafiltration assay (SITUA), was recently developed by the Nanotechnology Characterization Laboratory (NCL) [2, 15, 16]. This procedure addresses the major challenge of distinguishing the PK profiles of the drug fractions in plasma. A trace amount of isotopically labeled drug is used in the SITUA technique, and the assay assumes the drug isotope behaves identically to the original drug released from the nanomedicine in terms of protein binding [15]. That is, the drug isotope equilibrates between the unbound and protein-bound fractions. Subsequent separation by ultrafiltration separates the unbound drug (ultrafiltrate) from the encapsulated (nanomedicine-bound) and protein-bound drug fractions. The ultrafiltrate provides the measure of the unbound drug, from which the protein-bound and encapsulated fractions can be determined [16]. This technique enables the analysis of drug fractions derived from nanomedicines in the systemic circulation and can be easily adapted to a clinical setting. The SITUA method was recently used to analyze the PK profiles of Abraxane^®^ (nab-PTX), a clinically approved nanoformulation of the antineoplastic drug paclitaxel (PTX), and its nanosimilar formulation Genexol^®^ PM (nant-PTX) for bioequivalence testing in the Sprague-Dawley rat model [2]. The study revealed that despite having almost identical concentration-time profiles for total, unencapsulated, and unbound drug fractions, Abraxane^®^ and Genexol^®^ PM displayed some instances of statistically significant differences in the PK parameters, especially for the unencapsulated and unbound drug PK parameters, which was not apparent in the total drug PK profile [2]. These findings demonstrate the higher sensitivity of the SITUA approach in identifying modest deviations from bioequivalence that are not easy to detect with conventional techniques relying only on total drug PK.

We recently described a novel high-capacity polymeric micelle formulation of PTX in the triblock copolymer of (2-methyl-2-oxazoline) (PMeOx) and poly(2-butyl-2-oxazoline) (PBuOx) (PMeOx-*b*-PBuOx-*b*-PMeOx). This nanoformulated drug, termed POXOL *hl*-PM herein, has an unprecedented PTX loading capacity of up to 50% (wt.), thereby minimizing the total quantity of polymeric excipient required for solubilizing PTX [12]. As a result, potential excipient-related toxicity was reduced and the maximum tolerated dose (MTD) of POXOL *hl*-PM was increased compared to PTX (Taxol^®^) and Abraxane^®^ in mouse models [12]. In the current study, we present a preclinical evaluation of POXOL *hl*-PM and its bioequivalence in comparison to the clinically approved reference nanomedicine, Abraxane^®^. We utilized NCL’s SITUA method to assess the bioequivalence of the distinct subpopulations of POXOL *hl*-PM and Abraxane^®^ in two animal models (rats and rhesus macaques). The physicochemical properties, safety profile and antitumor efficacy of POXOL *hl*-PM in comparison with Abraxane^®^ are also presented. This study was carried out in collaboration with NCL, and the details of the process of the study are listed in Table 1.

**Table 1.**
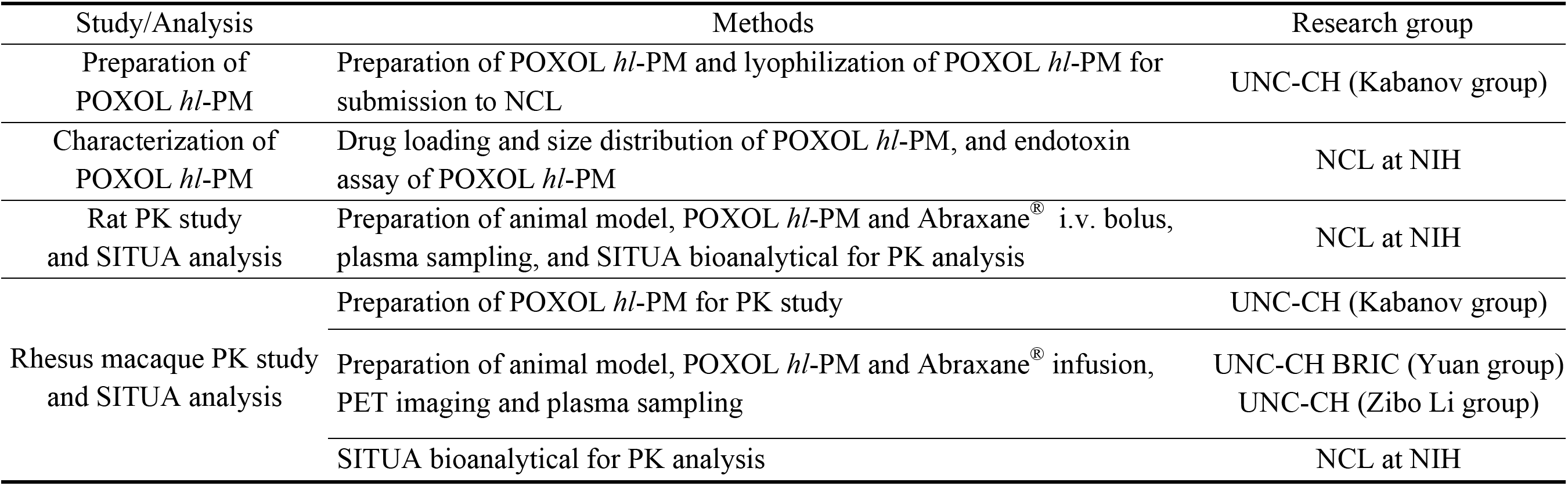
Summary of research workflow between the author groups/university research facilities and the NCL.

## 2. Materials and Methods

### 2.1 Materials

Amphiphilic triblock copolymer PMeOx_37_-*b*-PBuOx_21_-*b*-PMeOx_25_ (POx), (M_n_ = 8.1 kg/mol, D (M_w_/M_n_) = 1.27) was synthesized as described previously [17]. PTX was purchased from LC Laboratories (Woburn, MA). ^13^C_6_-Paclitaxel and ^2^H_5_-paclitaxel were purchased from Santa Cruz Biotechnology, Inc. (Dallas, TX). Abraxane^®^ is a protein-bound formulation of PTX manufactured by Celgene. 2-S-(4-isothiocyanatobenzyl)-1,4,7-triazacyclononae,1,4,7-triacetic (p-SCN-Bn-NOTA) acid was purchased from Macrocyclics (Plano, TX). All other materials were purchased either from Fisher Scientific Inc. (Fairlawn, NJ) or Millipore Sigma (St. Louis, MO). The MDA-MB-231 cell line was originally obtained from ATCC and cultured in Dulbecco’s modified eagle medium supplemented with 10% fetal bovine serum (FBS) and 1% penicillin-streptomycin. Matrigel was purchased from BD biosciences. ^64^Cu was obtained from the University of Wisconsin (Madison, WI). ^64^Cu was produced using the ^64^Ni (p, n) ^64^Cu nuclear reaction and supplied in high-specific activity as ^64^CuCl_2_ in 0.1 *N* HCl.

### 2.2 Preparation and Characterization of POXOL hl-PM

POXOL *hl*-PM formulations were prepared using a thin film method, as previously described [18]. In brief, a 4:10 weight ratio of PTX and POx polymer was dissolved in absolute ethanol and thoroughly mixed, followed by the complete removal of ethanol by a rotary evaporator. Sterile distilled water was used to rehydrate the films, and the micelle formulations were incubated at 60 °C for 20 minutes to form POXOL *hl*-PM solution. POXOL *hl*-PM solutions were transferred to sterile vials and freeze-dried for submission to NCL. Lyophilized powder form of POXOL *hl*-PM was reconstituted with sterile saline to prepare POXOL *hl*-PM solution. Physicochemical characterization of POXOL *hl*-PM was performed at NCL, including 1) PTX loading measurement by LC-Orbitrap system, 2) size distribution of POXOL *hl*-PM solution by dynamic light scattering (DLS), and 3) endotoxin study of POXOL *hl*-PM.

The amount of PTX encapsulated within the POXOL *hl*-PM was quantified with an LC-Orbitrap system (Thermo Fisher Scientific) using a Zorbax 300SB-C18 column (4.6 x 150 mm, 5 μm, 25 ^°^C). A gradient elution mode was used with a mobile phase composed of water with 0.1% trifluoroacetic acid (A) and acetonitrile with 0.1% trifluoroacetic acid (B). The gradient method consisted of the following segments: 50% B for 4 min, 10 min linear gradient to 100% B, hold at 100% B, 3 min linear gradient to 50% B, 50% B for 4 min. An injection volume of 40 μL (100-fold diluted) was used and a flow rate of the mobile phase was 1 mL/min. Loading efficiency (LE) and loading capacity (LC) of POXOL *hl*-PM were determined using the following equations (1–2): where M_PTX_ and M_POx_ are the weight amounts of the loaded (solubilized) PTX and POx in the POXOL *hl*-PM, while M_PTX added_ is the weight amount of PTX initially used for the preparation of POXOL *hl*-PM.

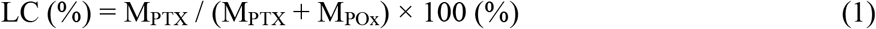

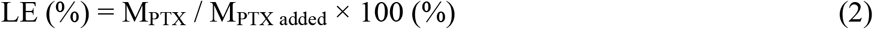

Size distribution of POXOL *hl*-PM solution was measured by DLS using Zetasizer Nano ZS (Malvern Instruments Ltd., UK). The POXOL *hl*-PM samples were diluted 10-fold and 100-fold in either water or saline, and the size distribution by intensity and volume were recorded, respectively. For the zeta potential measurement, the POXOL *hl*-PM samples were diluted 10-fold in water and brought to a PTX concentration of 0.2 mg/mL, and the apparent zeta potential was recorded at both the native and adjusted pH, at an applied voltage of 151 V.

### 2.3 In vitro endotoxin detection assay and assessment of the immunological response of POXOL hl-PM

The presence of Gram-negative bacterial endotoxin in the POXOL *hl*-PM micelles was tested using the kinetic Limulus Amebocyte Lysate assay protocol established by the NCL (https://ncl.cancer.gov/resources/assay-cascade-protocols). For the characterization of the immunological response of POXOL *hl*-PM, RAW264.7 mouse macrophage cell line cultured in DMEM supplemented with 10% heat-inactivated FBS in a 24 well plate was treated with the following: 1) lipopolysaccharide (positive control); 2) POXOL *hl*-PM; 3) POx; 4) DMSO-solubilized PTX. The cells were exposed to four concentrations of PTX (0.006, 0.03, 0.15, and 1.5 mg/mL). Twenty-four hours following treatment, the levels of TNF-α and MIP-1a were measured in the culture supernatant by enzyme-linked immunosorbent assay (R&D Systems) per the manufacturer’s instructions.

### 2.4 Husbandry

All animal experiments were performed in compliance with protocols approved by the local Institutional Care and Use Committee. The mice and rhesus macaques were housed in facilities accredited by the Association for the Assessment and Accreditation of Laboratory Animal Care International (AAALAC) at the University of North Carolina at Chapel Hill. Rats were housed at an AAALAC accredited facility in the Frederick National Laboratory for Cancer Research, and the experiments were conducted in accordance with the protocols approved by the NCI at Fredrick Institutional Animal Care and Use Committee.

Housing and husbandry for rhesus macaques strictly followed United States Department of Agriculture (USDA) animal welfare regulations. Rhesus macaques were housed in stable pairs and additional enrichments were provided in accordance with the standard operating procedure for non-human primate environmental enhancement to promote psychological well-being. All the study procedures were approved by the University of North Carolina at Chapel Hill Institutional Animal Care and Use Committee.

### 2.5 Determination of Toxicity and Efficacy of PTX formulations in mice

Tumor-free female nude mice (8 weeks old) were split into three groups (n=3) and given intravenous injections (q4d x 4 regimen) of one of the following treatments: 1) Normal saline; 2) POXOL *hl*-PM (125 mg/kg PTX equivalent); 3) POXOL *hl*-PM (150 mg/kg PTX equivalent). Four days after the final dose, mice were sacrificed, and the sciatic nerves were harvested using the mid-thigh incision technique [19]. Following formalin fixation, the tissue specimens were sectioned and stained with luxol fast blue combined with periodic acid-Schiff (LFB-PAS) for the histological examination of the morphology and myelination status of the peripheral nervous system following chemotherapy.

Female nude mice (6–8 weeks of age) were implanted with 5 x 10^4^ MDA-MB-231 breast cancer cells in 50% cell culture medium (FBS free) and 50% Matrigel (BD biosciences) by subcutaneous injection in the left dorsal flank. When tumors were about 100 mm^3^ in volume, tumor-bearing mice were randomized (n = 6 per group) and then administered with the following formulations: 1) sterile saline; 2) POXOL *hl*-PM (90 mg/kg PTX equivalent); 3) Abraxane^®^ (90 mg/kg PTX equivalent). The formulations were administered via tail vein following q4d x 4 regimen on days 0, 4, 8, and 12. Tumor growth was monitored twice weekly for 4 weeks. Tumor diameter ≥ 2 cm or moribund animals were set as the study endpoints. Tumor length (L) and width (W) were measured, and tumor volume (V) was calculated using the following equation: V = ½ x L x W^2^.

### 2.6 Pharmacokinetic studies in rat and rhesus macaques and toxicology study in rhesus macaques

Double jugular catheterized 15-week-old male Sprague Dawley rats were dosed with PTX formulations intravenously. Abraxane^®^ (12 mg/kg PTX equivalent) and POXOL *hl*-PM (12 mg/kg PTX equivalent) were injected via slow press bolus through the left jugular vein catheter (n = 7 per group). The right jugular vein catheter was used for blood sampling. 400 μL of blood was drawn at baseline, 15 min, 30 min, 1h, 2h, 4h, 6h, 7h, 8h, and 24h in K_2_EDTA tubes and spun at 2,500 x g at 37 °C for 10 min to separate the plasma for further analysis by SITUA, as described in section 2.8.

The PK study of the two formulations of PTX was also conducted in two female rhesus macaques (12 years age, ~8–9 kg) using a crossover design. The first monkey received an administration of POXOL *hl*-PM (12 mg/kg PTX equivalent), and Abraxane^®^ (12 mg/kg PTX equivalent) a month later, and the second monkey received POXOL *hl*-PM and Abraxane^®^ formulations in the reverse order. Administration of the two formulations was separated by at least 30-days of washout period to avoid any carryover. Both formulations were infused intravenously over 30 minutes. The blood samples were drawn before injection and at baseline, 5 min, 15 min, 20 min, 25 min, 30 min, 45 min, 60 min, 70 min, 100 min, 2 h, 24 h, and 48 h after injection in K_2_EDTA tubes and spun at 2,500 x g at 4 °C for 10 min and processed for further analysis by SITUA, as described in section 2.8.

One week following each injection of POXOL *hl*-PM and Abraxane^®^, blood was withdrawn and a comprehensive blood chemistry panel and complete blood count were performed for both the rhesus macaques. Blood was collected at one month following the second injection for the blood counts and chemistry analysis. Results of the blood analysis from two time points were compared.

### 2.7 Positron emission tomography/computed tomography imaging in rhesus macaque

^64^Cu-labeled POx was prepared for the *in vivo* biodistribution study of the polymer in rhesus macaque through positron emission tomography/computed tomography (PET/CT) imaging. The synthesis of POx with p-SCN-Bn-NOTA (NOTA-POx) was achieved by conjugating piperazine moiety to the end group of POx with p-SCN-Bn-NOTA as described previously [20]. Briefly, 20 mg of P(MeOx_37_-*b*-BuOx_21_-*b*-MeOx_25_) (2.47 μmol, 1 eq piperazine end group) was dissolved in 500 μL methanol along with 2.23 mg of p-SCN-Bn-NOTA (4.94 μmol, 2 eq), and approximately 5 mg anhydrous potassium carbonate was added. After incubating for 3 days at room temperature, the organic solvent was removed by evaporation, and the residual sample was collected with 1 mL of distilled water. NOTA-POx was purified from the residue by dialysis in a 3.5 kDa regenerated cellulose membrane in distilled water and lyophilized, following which NOTA-POx was obtained as a colorless solid.

For the chelation of ^64^Cu (half-life = 12.7 hours) to NOTA-POx, 2 mg of NOTA-POx and ^64^CuCl_2_ were dissolved in 200 μL of 0.1 *N* metal-free ammonium acetate buffer (pH 5.5). After 50 mins of incubation at 40 °C, the solution was allowed to cool down to room temperature. A small amount (10 μL) of 50 mM EDTA was added to the reaction mixture and unchelated radioactive copper ion was removed by centrifugation using Amicon^®^ Ultra centrifugal filters (regenerated cellulose, 3 kDa) (Merck Millipore, Billericia, MA). The radioactivity of ^64^Cu complexed NOTA-POx was measured in gamma counter prior to micelle preparation.

For imaging, the healthy rhesus macaque was anesthetized by isoflurane inhalation and placed on the imaging bed of a PET/CT system (Siemens Healthcare, Biograph64-mCT). Vital signs (ECG, pulse rate, blood pressure and body temperature) were monitored while the animal was under anesthesia. A whole-body CT scan was conducted for structure reference and attenuation correction for PET imaging. A single dose (~170 MBq) of ^64^Cu-labeled POXOL *hl*-PM was infused intravenously over 30 minutes with a syringe pump at a rate of 1 mL/min, which is a routine clinical practice for administering Abraxane^®^. Two-bed dynamic PET acquisition was started at the same time as the radiotracer infusion and continued for 1 hour. The animal was maintained under anesthesia for 3 hours, and the PET/CT scan was repeated at the 3 hours’ time point post-injection with static PET acquisition for 20 min. Following recovery from anesthesia, the animal was returned to the housing facility. Subsequently, repeated PET/CT scans were conducted at 24 and 48 hours post-injection to measure the biodistribution at later time points.

PET images were reconstructed using point spread function modeling and time-of-flight reconstruction algorithms with decay, scatter, and attenuation corrections. The 2-bed dynamic PET imaging acquired during the first hour after injection was assembled in 8-time frames (4 x 2.5 min, 3 x 5 min, 1 x 10 min) for each bed. Volumes of interest (VOIs) of major organs or tissue (heart ventricle, lung, liver, kidney, and muscle) were identified and demarked on the CT image and superimposed to the PET images. Time activity curves for the VOIs were obtained for up to 48 hours post-injection.

### 2.8 Stable isotope tracer ultrafiltration assay (SITUA)

The analysis of drug subpopulations in plasma was performed using the Nanotechnology Characterization Laboratory’s SITUA assay, as described previously [2, 16]. Briefly, plasma separated from blood samples was transferred to glass vials and spiked with ^13^C_6_-paclitaxel tracer at a concentration of 100 ng/mL. The spiked samples were vortexed and incubated at 37 °C with slight agitation for 10 min. A 25 μL volume of the spiked sample was removed and treated with acetonitrile (0.1% formic acid) for protein precipitation and used for the analysis of total drug concentration. The remainder of the plasma was subjected to ultracentrifugation in a 10 kDa molecular weight cut-off filter (Vivacon) at 12,000 g for 10 min at 37 °C. The ultrafiltrate was used for determining the percentage of the unbound drug following protein precipitation. A 25 ng/mL concentration of ^2^H_5_-paclitaxel was used as an internal standard for every sample. The samples were stored at –80 °C until further analysis. On the day of the analysis, the samples were thawed and dried in a centrifuge speed vacuum and reconstituted with 40% acetonitrile/0.1% formic acid, and analyzed on a Q – Orbitrap. Thereafter, the encapsulated, protein-bound, free drug fractions were determined as described previously [2].

To assess the effect of the freeze-thaw cycle on the release of PTX from either POXOL *hl*-PM or Abraxane^®^, the two formulations were spiked into rhesus macaque plasma at 5 μg/mL PTX equivalent in triplicate. A portion of the freshly spiked plasma was analyzed by SITUA, and the remainder was frozen at −80 °C for 24 hours and subjected to SITUA the next day.

### 2.10 Pharmacokinetic data analysis

PK data analysis was performed using protocols published by NCL, as previously described [2]. Briefly, the PK parameters were calculated by noncompartmental analysis using Phoenix WinNonlin 8.2 software (Pharsight Corporation, Mountain View, CA). The first-order trapezoidal rule was used to calculate the area under the plasma concentration/time curve from time zero to 48 hours (AUC_all_) and estimate the area under the curve to time infinity (AUC_inf_) by extrapolating AUC to time infinity. The maximum drug concentration (C_max_) and the time of maximum concentration (T_max_) were obtained from the measured data. Clearance (CL), volume of distribution steady state (V_ss_), mean residence time (MRT_inf_) and half-life (T_1/2_) were calculated using equations (3), (4), (5), and (6), respectively; λ_z_ represents the natural log slope of terminal elimination.

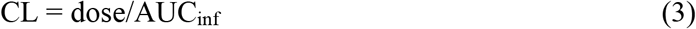

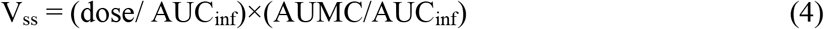

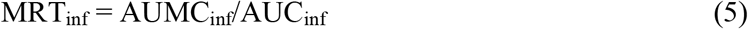

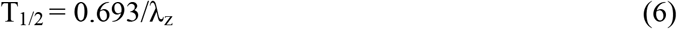

### 2.11 Statistics

Statistical differences were determined by Student’s *t*-test or one-way ANOVA with Tukey’s posthoc test for multiple comparisons, with a significance level, α, set at 0.05.

## 3. Results

### 3.1 POXOL *hl*-PM *formulation*

Three independent batches of PTX polymeric micelle formulation (POXOL *hl*-PM) were manufactured at laboratory scale (see **Supplementary Table S1**) for preclinical evaluation using the thin film technique as previously described [18]. Batches 2 and 3 of POXOL *hl*-PM contain 6.9 mg PTX per 16.0 mg of POx and represent spherical micelles with a particle size of 30 nm and polydispersity index (PDI) of 0.05, as determined by DLS (**Supplementary Table S2**). Particle distribution of POXOL *hl*-PM was similar to the PTX-loaded POx micelle formulation described previously [12], albeit the POx used in this formulation had a shorter PMeOx block (**Supplementary Table S1**). Additional characterization data on PTX loading in POXOL *hl*-PM and size distribution upon dilution is presented in Supplementary materials (**Supplementary Fig. S1**, **Table S2**). POXOL *hl*-PM displayed a concentration-dependent size distribution, with narrower size distributions apparent at lower dilutions. This was true for both micelles dispersed in saline as well as DI water (**Supplementary Fig. S1**). Also, the micelles exhibited a neutral zeta potential when dispersed in water and PTX loading of about 26% at a POx/PTX weight ratio of 10/4 (**Supplementary Table S2**).

The induction of proinflammatory cytokines from RAW264.7 cells was studied to evaluate the presence, if any, of proinflammatory effects of POXOL *hl*-PM. The treatment of RAW264.7 cells with POXOL *hl*-PM (batch 1) induced TNF-α and MIP-1a for all the concentrations tested, similar to the DMSO-solubilized PTX (**Supplementary Fig. S2**). The immunogenic response elicited by PTX is due to its pharmacological activity and is consistent with previous reports [21]. In contrast, POx alone did not induce either cytokine at any of the tested concentrations suggesting that the induction of the cytokines by POXOL *hl*-PM was solely due to PTX. These results, together with the previously reported negligible complement stimulation activity of POx, exemplify the non-immunogenicity of POx polymers [12].

### 3.2 Safety and anti-tumor effect of POXOL hl-PM in mouse models

The maximum tolerated dose (MTD) for POXOL *hl*-PM was determined in healthy 6–8 weeks old female nude mice using a q4d x 4 regimen. The body weight changes were monitored during the treatment, and the mice group injected with POXOL *hl*-PM showed significant body weight loss (> 15%) at a PTX dose of 150 mg/kg. When treated with 125 mg/kg dose of PTX, the bodyweight loss was less than 15% (**Fig. 1A**). It was thus concluded that the MTD for POXOL *hl*-PM determined under q4d x4 regimen was 125 mg/kg of PTX by POXOL *hl*-PM, which was similar to the previously reported MTD of 150 mg/kg of PTX for PTX-loaded POx micelles in mouse [12]. Differences in MTD between the two studies might have been due to minor changes in the formulation, as we realized that the block lengths of the hydrophilic repeat segments in the POx used in the present study differed from that of the polymer used previously, presumably resulting in a lower hydrophilic/lipophilic balance of the POx due to a relatively shorter hydrophilic block in the present study. The difference in the hydrophilic/lipophilic balance could result in differences in the retention of PTX by the two polymeric micelles, thereby affecting drug disposition and subsequent drug toxicity profile and weight loss. Abraxane^®^ was used in this study as the reference drug. We previously determined its MTD to be 90 mg/kg in the same dosage regimen, in the same strain of mice [12].

**Fig. 1.**
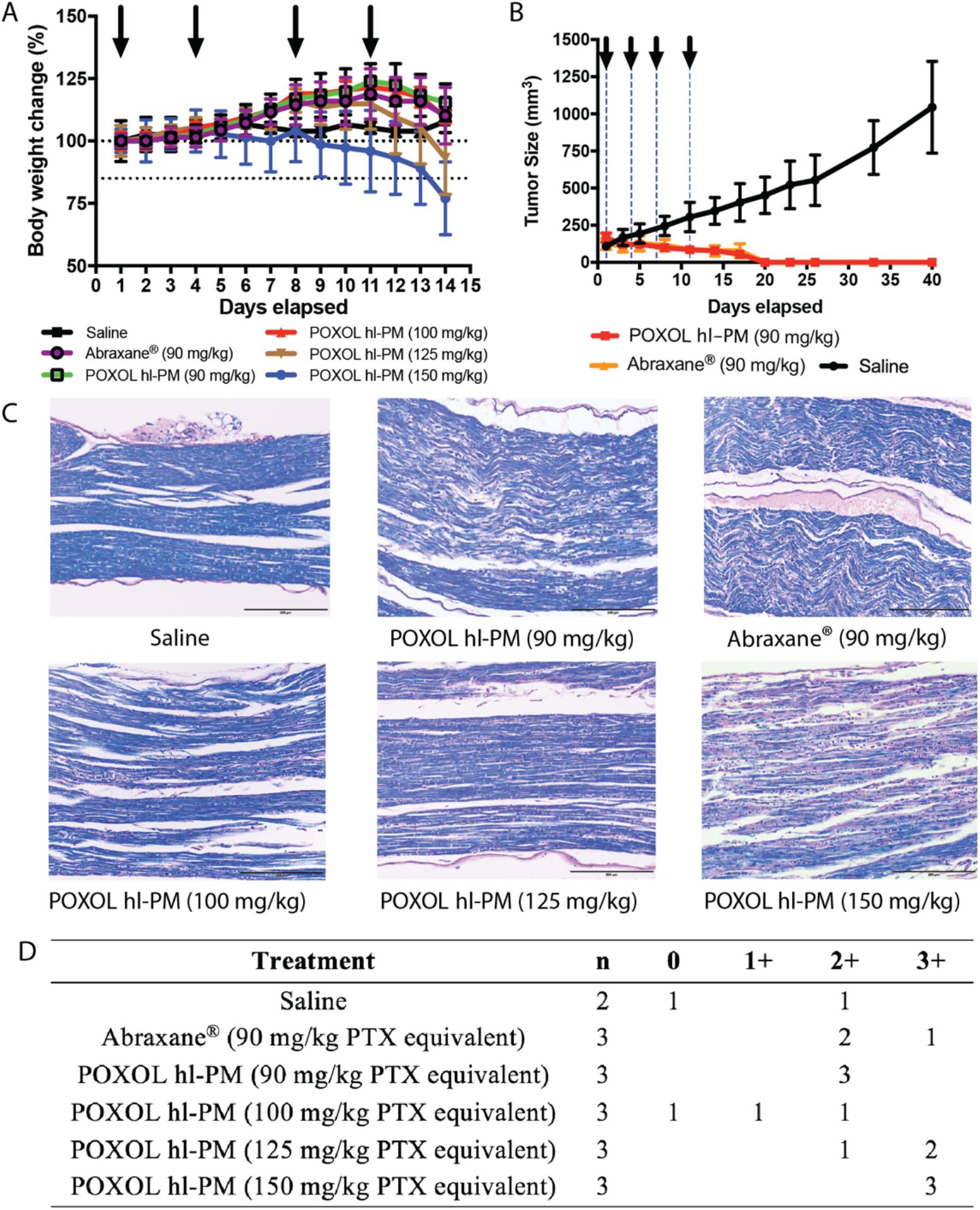
Toxicity and efficacy assessment of POXOL *hl*-PM and Abraxane^®^ in mouse. **(A)** MTD of PTX equivalent (batch 2) **(B)** Tumor inhibition in MDA-MB-231 human breast xenograft model. (**C**) LFB-PAS-stained sciatic nerve sections for the histological assessment of neurotoxicity following treatments with saline, Abraxane^®^ (90 mg/kg PTX equivalent) and POXOL hl-PM (90 mg/kg, 125 mg/kg, and 150 mg/kg PTX equivalent). **(D)** Scoring of neuropathy in a 4-point scale.

A tumor inhibition study was performed to evaluate the antitumor efficacy of POXOL *hl*-PM and Abraxane^®^ in the MDA-MB-231 xenograft model. Tumor-bearing mice were treated with POXOL *hl*-PM or Abraxane ^®^ at a 90 mg/kg PTX equivalent (q4d x 4 regimen). Compared to the control group (treated with sterile saline), both POXOL *hl*-PM and Abraxane^®^ treated groups showed complete remission of the tumor, and tumor regrowth was not observed when examined for an additional 25 days (**Fig. 1B**). In contrast, the tumors in the control group attained an average size of 1,000 mm^3^ by day 40 (study termination). The tumor inhibition study revealed that both POXOL *hl*-PM and Abraxane^®^ had comparable antitumor efficacy in the xenograft tumor model.

Neurotoxicity is a well-documented side effect of PTX in both mice and humans [22, 23]. Demyelination of nerve fibers is secondary to axonal degeneration [24], and therefore, we sought to characterize PTX neuropathy by studying the demyelination of the sciatic nerves in mice treated with POXOL *hl*-PM or Abraxane^®^ (q4d x 4 regimen). To this end, LFB-PAS stain was used, which differentiates myelinated regions (blue stain) from demyelinated regions (pink stain). A 4-point scoring system (neg–no degenerative changes; 1-slight degree of degenerative changes; 2-moderate degenerative changes; 3-marked degree of degenerative changes) was employed to assess the degree of neuropathy. Abraxane^®^ displayed comparable severity of neuropathy (avg. score: 2.2) to POXOL *hl*-PM (avg. score: 2) at the same dose (**Fig. 1C, D**). While one of the control samples displayed a moderate level of degeneration, this could have been an artifact of the isolation procedure because the mice in this group received only saline. At higher doses of POXOL *hl*-PM (100, 125, and 150 mg/kg PTX equivalent), the mice displayed a dose-dependent neurotoxic response to PTX, with a higher degree of nerve degeneration observed at higher doses (**Fig. 1C, D**).

### 3.3 Pharmacokinetic study in rats and rhesus macaques and toxicology study in rhesus macaques

Two animal models (rats and rhesus macaque) were employed to investigate the PK profiles of drug fractions from POXOL hl-PM and Abraxane^®^ using the SITUA method described above (**Fig. 2A**). The 12 mg/kg PTX equivalent dose used for the PK studies was derived from dose conversion of the highest tolerated clinical dose of the clinically approved PTX polymeric micelle product (Genexol^®^PM (435 mg/m^2^), Cynviloq (USA)) (approved in South Korea, Philippines, Vietnam, and India) [25]. The mg/kg PTX equivalent dose was used directly without body surface area conversion since the volume of distribution has an interspecies allometric scaling exponent of 1 (i.e., V_d_ change is directly proportional to body weight). (see **Supplementary Table S3** and supplementary materials for additional information on dose conversion). Blood samples were obtained at predetermined time points, and drug fractions from POXOL hl-PM and Abraxane^®^ were analyzed by SITUA (**Fig. 2A**). The PK profiles (total, encapsulated, protein-bound, and free PTX) were analyzed to evaluate the bioequivalence of POXOL *hl*-PM and Abraxane^®^ in given animal models.

**Fig. 2.**
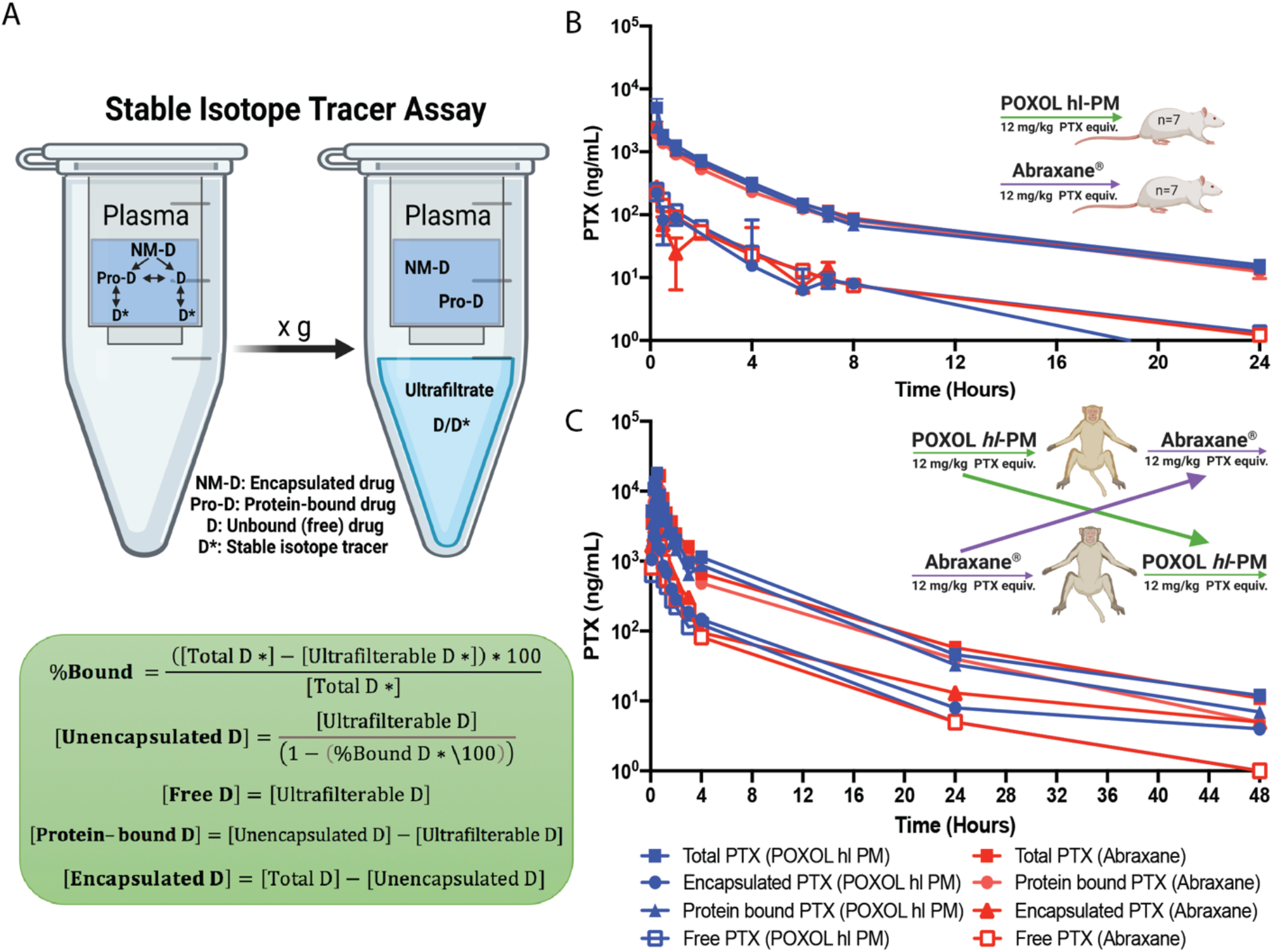
Pharmacokinetics of POXOL and Abraxane^®^ in rats and rhesus macaques. **(A)** Schematic illustration of stable isotope ultrafiltration assay (SITUA) (created with BioRender.com). Equations for free, encapsulated and protein-bound PTX time-concentration comparison for POXOL (batch 3) and Abraxane^®^ in **(B**) utilized rats and **(C)** rhesus macaques. A two-arm parallel study design was in rats (n=7-8) and a crossover design was used in rhesus macaques (n=2). (**B and C**)

In rats, both Abraxane^®^ and POXOL *hl*-PM displayed triphasic concentration-time curves for all the drug populations (**Fig. 2B**). The free drug represented approximately 10 % of the total drug. The total drug concentration was dominated by the protein-bound drug and was almost identical between Abraxane^®^ and POXOL *hl*-PM (**Table 1** and **Supplementary Table S4**). The encapsulated drug was present in a low quantity for both formulations. The PK parameters for Abraxane^®^ and POXOL *hl*-PM were found not to be statistically different by Student’s t-test (p<0.05). These results suggest that in addition to the total drug PK, subset drug fractions such as encapsulated, protein-bound, and unbound drug PK were bioequivalent in rats.

**Table 1.** Comparison of pharmacokinetic parameters of POXOL hl-PM and Abraxane^®^ in rats.

| Parameters<sup>a</sup> | POXOL *hl*-PM |  |  |  | Abraxane<sup>®</sup> |  |  |  |
| --- | --- | --- | --- | --- | --- | --- | --- | --- |
|  | Total | Protein-bound | Encapsulated | Free | Total | Protein-bound | Encapsulated | Free |
| T<sub>max</sub> (h) | 0.3 | 0.3 | 0.5 | 0.3 | 0.3 | 0.3 | 0.3 | 0.3 |
| C<sub>max</sub> (ng/mL) | 2915.9 | 2441.9 | 292.7 | 260.3 | 2418.5 | 1931.8 | 289.5 | 206.5 |
| AUC<sub>all</sub> (ng•h/mL) | 5600.1 | 4787.5 | 299.7 | 485.3 | 4864.5 | 4089.7 | 465.4 | 414.7 |
| AUC<sub>inf</sub> (ng•h/mL) | 5755.2 | 4936.2 | 327.7 | 498.8 | 4978.8 | 4197.9 | 489.6 | 425.8 |
| CL (mL/h•kg) | 2185.3 | 2539.5 | 14785.2 | 25348.1 | 2562.7 | 3036.8 | 66001.4 | 29959.8 |
| t<sub>1/2</sub> (h) | 6.4 | 6.7 | 5.5 | 6.5 | 5.8 | 5.7 | 5.5 | 5.8 |
| MRT (h) | 4.2 | 4.3 | 4.5 | 4.3 | 4.1 | 4.2 | 2.7 | 4.1 |
| V<sub>ss</sub> (mL/kg) | 9530.7 | 11502.3 | 114972.3 | 114060.3 | 10597.1 | 12718.7 | 156655.9 | 118992.3 |

<sup>a</sup> T<sub>max</sub>, time of maximum concentration; C<sub>max</sub>, maximum plasma concentration; AUC<sub>all</sub>, area under the curve from all time points; AUC<sub>inf</sub>, area under the curve until infinite time; CL, clearance; t<sub>1/2</sub>, half life; MRT, mean residence time; V<sub>ss</sub>, apparent volume of distribution at steady state; Data represents mean of 7 animals; data for individual animals can be found in **Table S4**.

In rhesus macaques, plasma analysis by the SITUA method revealed that PK profiles of the total, protein-bound, encapsulated, and free PTX were similar, and the calculated PK parameters were not significantly different between the two formulations (confirmed by Student’s t-test) (**Table 2** and **Supplementary Table S5)**. Free drug represents approximately 10% of the total drug, leaving the rest (and majority) as protein bound (**Supplementary Table S5**). The apparent nanomedicine encapsulated fraction represented approximately 15-20% of the total drug, although this fraction could be misrepresented by our assay as a result of the limitations of the study as discussed below. These results indicate that the drug partitioning from Abraxane^®^ and POXOL hl-PM to plasma albumin occurs to a similar extent in systemic circulation. This data supports the bioequivalence of POXOL hl-PM and Abraxane^®^ in rhesus macaques.

**Table 2.** Comparison of pharmacokinetic parameters of POXOL hl-PM and Abraxane^®^ in rhesus macaque.

| Parameters<sup>a</sup> | POXOL *hl*-PM |  |  |  | Abraxane<sup>®</sup> |  |  |  | POx<sup>b</sup> |
| --- | --- | --- | --- | --- | --- | --- | --- | --- | --- |
|  | Total | Protein-bound | Encapsulated | Free | Total | Protein-bound | Encapsulated | Free | Total |
| T<sub>max</sub> (h) | 0.6 | 0.6 | 0.6 | 0.6 | 0.5 | 0.5 | 0.5 | 0.5 | 0.58 |
| C<sub>max</sub> (ng/mL) | 14944.0 | 9586.5 | 4301.5 | 1649.0 | 17899.5 | 11956.7 | 4127.0 | 1815.8 | 296602.6 |
| AUC<sub>all</sub> (ng•h/mL) | 22198.2 | 15072.5 | 4835.9 | 2567.6 | 22276.2 | 15801.8 | 3991.1 | 2460.5 | 2797225.9 |
| AUC<sub>inf</sub> (ng•h/mL) | 22319.6 | 15123.9 | 4918.7 | 2574.7 | 22440.2 | 15874.3 | 4032.3 | 2474.5 | 2837613.5 |
| CL (mL/h•kg) | 542.4 | 798.7 | 2440.7 | 4793.4 | 546.1 | 776.6 | 2999.6 | 4935.0 | 10.57 |
| t<sub>1/2</sub> (h) | 7.6 | 6.8 | 10.7 | 6.6 | 9.3 | 6.8 | 8.1 | 8.8 | 7.51 |
| MRT (h) | 4.0 | 34.0 | 3.8 | 3.5 | 4.1 | 4.1 | 4.0 | 4.0 | 9.8 |
| V<sub>ss</sub> (mL/kg) | 2310.9 | 3286.6 | 11890.8 | 17440.1 | 2552.4 | 3486.1 | 13896.4 | 21670.9 | 110.99 |

<sup>a</sup> T<sub>max</sub>, time of maximum concentration; C<sub>max</sub>, maximum plasma concentration; AUC<sub>all</sub>, area under the curve from all time points; AUC<sub>inf</sub>, area under the curve until infinite time; CL, clearance; t<sub>1/2</sub>, half life; MRT, mean residence time; V<sub>ss</sub>, apparent volume of distribution at steady state; <sup>b</sup> PK parameter of POx was determined from the PET/CT study described in section Result (section 3.4). Data represents mean of 2 animals; data for individual animals can be found in **Table S5**.

Since the plasma samples had to be frozen before being shipped to NCL for SITUA analysis, a freeze-thaw control experiment was carried out. The study examined the effect of freeze-thaw procedure on the composition of PTX subpopulations in the plasma, to ensure that the procedure did not result in the premature release of PTX from either formulation. Both fresh and frozen samples yielded comparable percentages of encapsulated and unencapsulated drug for both formulations (**Supplementary Table S6**). Therefore, the freezing procedure did not alter the extent of PTX release from either POXOL hl-PM or Abraxane^®^.

Blood samples were collected at one week and one month after the administration of POXOL *hl*-PM_or Abraxane^®^ to conduct complete blood counts, and comprehensive metabolic panel to assess the toxicity of the formulation. As demonstrated in **Supplementary Table S7**, POXOL *hl*-PM induced a much lower change in monocyte, lymphocyte and platelet counts compared to Abraxane^®^. This can be attributed to the development of hypersensitivity reaction to human serum albumin (HSA) present in Abraxane^®^, which has been previously reported in rhesus macaques and dogs infused with HSA [26, 27]. Furthermore, some acute side effect symptoms were reported by the veterinarian after treatment with Abraxane^®^, but not after POXOL hl-PM. These side effect symptoms included ataxia, ventricular premature contractions (VPCs) every 2-3 minutes on the EKG about 45mins-1h post-Abraxane^®^ administration, protracted course of diarrhea, and diffuse alopecia of forelimbs, hindlimbs, and back. As for CMP, although there was a slight elevation of alkaline phosphatase (ALK), an indicator of liver function/toxicity, the level was still within the normal range (**Supplementary Table S8**). The overall toxicity analysis demonstrated a comparable blood toxicity profile of POXOL *hl*-PM to that of Abraxane^®^ at the given dose of PTX in rhesus macaques.

### 3.4 PET-CT Biodistribution study of 64Cu-labeled POXOL hl-PM in rhesus macaques

We conducted a pilot Positron Emission Tomography – Computed Tomography (PET/CT) imaging study to determine the biodistribution and PK of POXOL hl-PM in a naive rhesus macaque model. As shown in **Fig. 3A**. Time activity curves for each organ were obtained to assess the tissue PK of ^64^Cu-labeled POx. Initially, the highest activity of ^64^Cu-labeled POx was detected in the blood, which was demonstrated by high radioactivity in the heart ventricle. In the kidney, we observed a strong signal at initial time points (during infusion), following which the signal gradually declined (**Fig. 3A**). The liver and spleen were other organs where the high activities of ^64^Cu-labeled POx were detected. Over time, all signals gradually declined. The signal from blood and kidney declined faster, while sustained signals from the liver and spleen were observed. In the case of muscle, we observed a marginal signal of ^64^Cu-labeled POx, indicating that POx does not accumulate in the muscle (**Fig. 3A**). The PET-CT images of ^64^Cu-labeled POx in rhesus macaques depicted that POXOL hl-PM could circulate in the body for up to 48 hours in rhesus macaques (**Fig. 3B**).

**Fig. 3.**
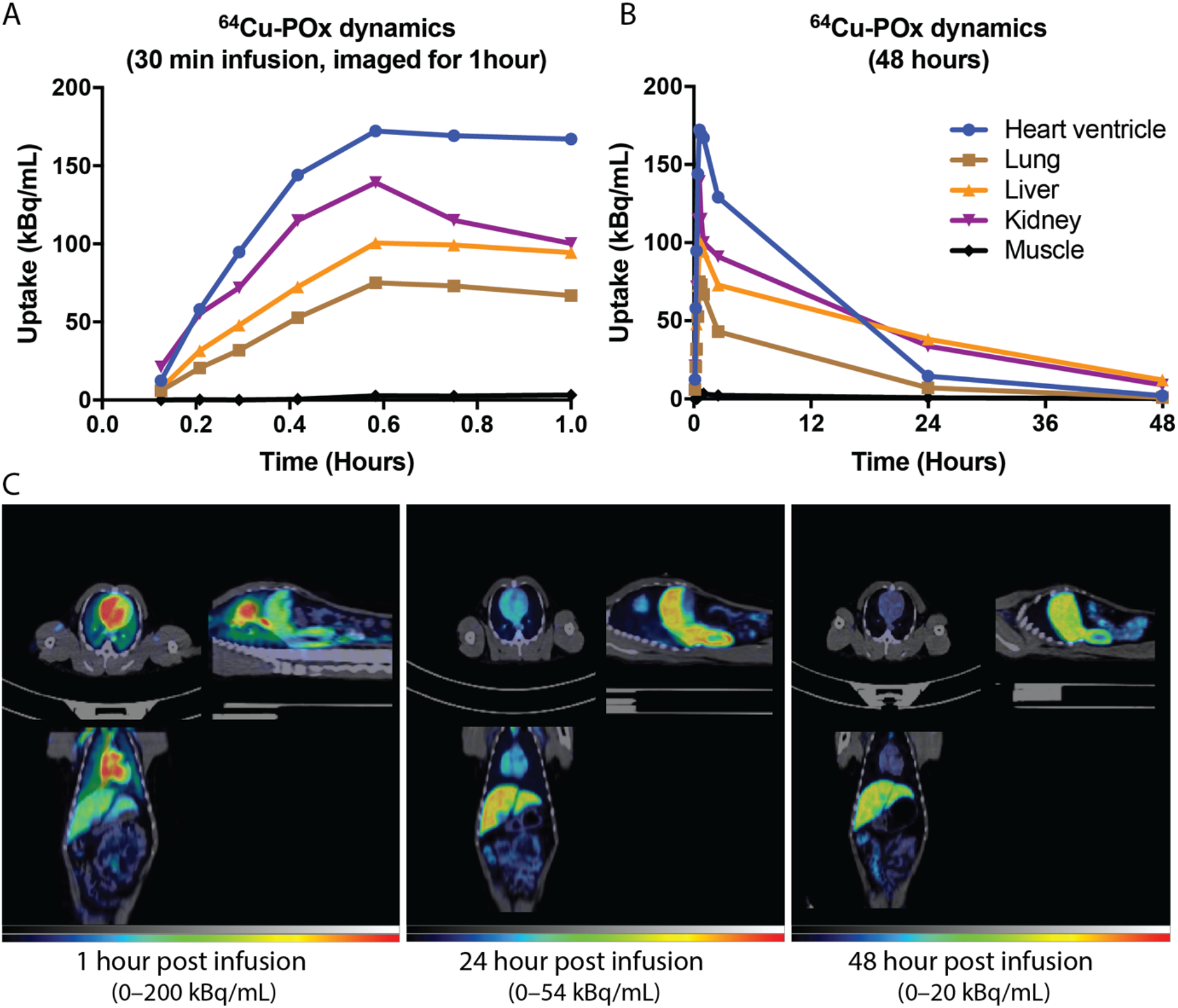
POx biodistribution in rhesus macaque. Presented are the results of the tissue biodistribution of ^64^Cu-labeled POx for **(A)** 1 h and **(B)** 40 h post infusion of ^64^Cu-labeled POXOL hl-PM. **(C)** PET/CT images in the (clockwise from upper left) transverse, sagittal, and coronal plane of the rhesus macaque at different time points following ^64^Cu-labeled POXOL hl-PM infusion.

Most notably, the POx polymer has a much lower volume of distribution (roughly 25-fold) than the drug itself (**Table 2**). This indicates that while the drug readily penetrates peripheral tissues, the polymer cannot achieve this to the same extent. The half-life for all fractions ranged from 6.75 to 8.14 hours for POXOL hl-PM and 6.58 to 10.24 hours for Abraxane^®^. The clearance values for the drug fractions in each formulation were remarkably similar across formulations. The clearance of the polymer is roughly 50-fold less than that of the total drug in each formulation (**Table 2**).

## Discussion

Emerging evidence of improved safety and efficacy of nanoformulated drugs compared to conventional drug formats has motivated the increased development of nanoparticle-based drug delivery systems over the last few decades [4, 28, 29]. Nonetheless, the clinical translation of nanoformulated drugs has been relatively slow [28]. One potential reason for this could be the lack of comprehensive preclinical PK evaluation methodologies, limiting the ability to accurately determine the altered PK/PD relationship of nanoformulated drugs. For instance, most PK studies rely on the measurement of the total drug and do not focus on the relative concentrations of nanoparticle encapsulated, protein-bound, and free drugs released from the nanocarrier in the systemic circulation. However, reliance on the total drug amount in the plasma following intravenous administration of nanoformulated drugs may inaccurately estimate drug exposure as it doesn’t account for the nanomedicine-derived drug subpopulations [3]. Such subpopulations are dynamically related to each other and able to alter the PK/PD relationship of the drug and ultimately affect therapeutic outcomes [12, 13]. Thus, the presence of nanocarrier-associated drugs during circulation and subsequent drug release from nanomedicine adds an additional layer of complexity to evaluation of drug disposition. To decipher this complex and dynamic drug PK profile composed of drug subpopulations from nanomedicine, robust analytical techniques are needed.

The altered PK profile of nanoformulated drugs compared to conventional drug formats could influence the tissue accumulation and toxicity profile of the nanomedicine as seen in preclinical studies and clinical trials. For instance, the non-linear PK of Taxol^®^ attributed to the equilibrium binding of the drug to the micelle-forming surfactant excipient, Cremophor-EL can result in a disproportionate increase in total drug AUC with increasing drug dose [30]. The analysis of the unbound drug in PK models revealed that the dose-dependent PK of Taxol^®^ may have clinical consequence because it could result in a decrease in the unbound PTX fraction and lower the tissue distribution and clearance of the drug. A study in ovarian cancer patients showed that a 30% increase (from 135 mg/m^2^ to 175 mg/m^2^ PTX) in the dose of Taxol^®^ (administered through 3h infusion) resulted in a 75% increase in the mean AUC of PTX [30]. The ramifications of this effect could also extend to the off-target effects of Taxol^®^.

In addition to the toxicity of the PTX molecule, Taxol^®^ has off-target effects such as Cremophor-EL-mediated complement system activation and hypersensitivity reactions resulting from Cremophor-EL-induced histamine release, which limit the therapeutic window of Taxol^®^ [31]. By replacing Cremophor EL with a safer excipient (HSA), Abraxane^®^ (nab-PTX) was able to alleviate the dose-limiting toxicity of Taxol^®^. Unlike Taxol^®^, Abraxane^®^ exhibits linear PK behavior that produces a proportionate dose-exposure relationship [32]. Therefore, despite having the same active pharmaceutical ingredient, Taxol^®^ (PTX formulated in Cremophor EL and ethanol) has a much lower MTD than Abraxane^®^ (<50%), which has been a major impediment to its clinical utility by limiting its use to lower doses of PTX in patients [7]. Genexol^®^PM (Nant-PTX) is a Cremophor EL-free polymeric micelle formulation of PTX based on poly(ethylene glycol)-*b*-poly(D,L-lactide) block copolymer. Genexol^®^ PM also has a linear PK profile in humans with an MTD of 435 mg/m^2^, implying enhanced safety of this drug nanoformulation [25]. Thus, it is understood that the PK and toxicity of a nanoformulated drug is dependent upon both the carrier (excipient) and the drug, and the carrier can dramatically influence drug disposition and tissue exposure.

The NCL was established to support preclinical analyses of nanoformulated drugs and devices, and streamline efforts at clinical translation. The NCL offers a gamut of standardized analytical protocols to evaluate the physicochemical properties of the nanoformulated drugs and assess their toxicity profiles both *in vitro* and *in vivo* (https://ncl.cancer.gov/resources/assay-cascade-protocols). In the present study, in collaboration with NCL, we conducted a comprehensive analysis of POXOL hl-PM to better understand the properties of the nanoformulation prior to clinical translation. This study includes characterization of drug loading and particle size distribution, endotoxin assay, and advanced PK analysis and bioequivalence evaluation of POXOL hl-PM in various animal models using the established SITUA bioanalytical method. This is an example of how the well-designed infrastructure for nanomedicine characterization at NCL can be utilized by nanoformulation scientists to improve the prospects of clinical translation (https://ncl.cancer.gov/working-ncl/ncl-assay-cascade-application-process).

POXOL *hl*-PM is unique in that it uses significantly less excipient than other drug delivery platforms due to POx polymeric micelles’ high solubilizing capacity for poorly soluble drugs, thereby improving safety substantially [12, 33, 34]. We have previously demonstrated that for poorly water-soluble drugs such as PTX, concentrations as high as 50 g/L are attainable by POx polymeric micelles, which is ten orders of magnitude higher than Abraxane^®^ (5 g/L) [12]. The studies undertaken to understand the high PTX-loading capacity of POx micelles identified two distinct molecular mechanisms of PTX solubilization by POx: 1) the formation of a distinct microarchitecture in the core following PTX encapsulation by POx micelles [35, 36] and 2) hydrophilic block (PMeOx)-assisted PTX solubilization in POx micelles at high capacity [37].

In addition to its unique solubilization property, POx has also been shown to have long shelf stability and be hematologically and immunologically safe in a rodent (mouse) model [12]. In the current study, we compared the antitumor efficacy of POXOL hl-PM and Abraxane^®^ dosed at the MTD of the latter (90 mg/kg PTX equivalent) in the MDA-MB-231 mouse model of breast cancer. Our study revealed that POXOL hl-PM was therapeutically comparable to Abraxane^®^ with regard to tumor inhibition activity at the same dose of PTX and regimen. We further investigated the immunostimulatory effect of POXOL hl-PM and POx to assess the biocompatibility of the polymer. The lack of inflammatory activity of polymer in RAW-264 macrophages demonstrated in this study further corroborates the non-immunogenic nature of the POx as an excipient of POXOL hl-PM.

Since demyelination plays a critical role in the pathophysiology of PTX neuropathy [24], we sought to assess the degree of demyelination in sciatic nerves in mice following treatment with four systemic doses of Abraxane^®^ or POXOL hl-PM. In our study the extent of demyelination caused by Abraxane^®^ was comparable to, if not higher than, that of POXOL hl-PM. The dose-dependent severity of PTX-induced neuropathy has been reported in clinical studies of Abraxane^®^ and Taxol^®^ [38]. Abraxane^®^ dosed at 260 mg/m^2^ (q3w) produced grade 3 sensory neuropathy in 10% of the breast cancer patients compared to the 2% with Taxol^®^ at 175 mg/m^2^ (q3w) [32]. However, the incidence of grade 3 neuropathy in the Abraxane^®^ arm was still lower than that observed with Taxol^®^ (32 %) at a dose of 250 mg/m^2^ in a separate study using the same dosing schedule. This could have been because of the cumulative toxicity of PTX and Cremophor EL since Cremophor EL also contributes to neuropathy. It is evident from these instances that Cremophor EL-free formulations of PTX have more favorable toxicity profiles.

The ability to distinguish between individual drug subpopulations in circulation makes SITUA an ideal tool for characterizing nanoformulated drugs. The assay revealed that the drug was present in three forms in the systemic circulation, including micelle encapsulated, protein-bound, and unbound. As expected from previous reports, the protein-bound fraction was the most prevalent of the three [2]. The percentage of unencapsulated or released drug was comparable for the Abraxane^®^ and POXOL hl-PM formulations at all time points measured in both models, implying equivalent release rates. This value remained relatively constant at 80% in the rhesus macaque bioequivalence study, indicating that 20% of the total drug in circulation was retained in the formulation for the first 24 hours. In the rat bioequivalence study, the encapsulated drug was detectable in the early time points for both POXOL hl-PM and Abraxane^®^, with nearly 5% of the drug retained in the formulation in the first 1 hour. By contrast, the SITUA study performed previously for assessing the bioequivalence of Abraxane^®^ and Genexol^®^PM in rats reported undetectable encapsulated drug even at the earliest time point for the Abraxane^®^ group. This is an important finding as the dose of PTX used for that study (6 mg/kg) was the half of the dose used in the current study (12 mg/kg), raising the question of the effect of dose on the release and, therefore the stability of Abraxane^®^ nanomedicine. This phenomenon cannot be attributed to saturated protein binding since the dose used in the current study is not high enough to produce the saturation of drug-protein binding *in vivo*. More research into the relationship between dose and stability of nanomedicine is thus required. It is our opinion, that once the drug is injected into the body as a micelle, it rapidly binds to the sink of proteins in circulation, causing the majority of the drug to be released rapidly from the formulation. However, a significant portion of the drug remains encapsulated in the micelle, which could play a critical role in PK activity (influence the altered PK/PD) as this could provide alternative mechanisms of uptake into sites of therapeutic interest, like tumors.

This study has limitations, particularly with the estimation of the encapsulated fraction from the SITUA. Since the encapsulated fraction is obtained from the difference of the measure of the total drug (obtained directly from the reservoir) and the unencapsulated fraction estimated using the drug isotope, any error in the estimation of the unencapsulated fraction can be carried forward in the estimation of the encapsulated fraction. As a result, the unencapsulated drug estimate may be ± error (%) of the nominal concentration of the unencapsulated drug. The FDA guidance document on bioanalytical method validation states that the acceptance limit for accuracy for chromatographic assays should be ± 20% of the nominal concentration (https://www.fda.gov/files/drugs/published/Bioanalytical-Method-Validation-Guidance-for-Industry.pdf), and the accuracy deviation for SITUA is within this acceptance criterion [2]. In addition, the freeze-thaw cycle could introduce additional errors in the measurement of the drug. For instance, in the present study, there was 90% release of PTX in the freshly prepared samples, compared to the 80% release of PTX in the frozen samples. Although the difference was not statistically significant, the freeze/thaw process could skew the estimation of drug fractions from SITUA. Therefore, further validation studies for SITUA are needed to incorporate the dynamic nature of drug fractions from polymeric micelle formulation and account for the changes in experimental conditions.

There is preliminary evidence for the bioequivalence of Genexol^®^PM to Abraxane^®^. Their bioequivalence was revealed by the data collected from eight patients in a 2014 clinical trial study comprising patients with advanced breast cancer and non-small cell lung cancer (NSCLC) patients (NCT02064829). However, the bioequivalence study only compared the total PTX and did not investigate the PTX subpopulations. Recent FDA guidance documents for drug products containing nanomaterials states the need to measure all drug fractions for establishing bioequivalence [3]. In the present study, the bioequivalence of Abraxane^®^ and POXOL hl-PM in rats and rhesus macaques was investigated for all subpopulations of PTX using SITUA. The bioequivalence of POXOL hl-PM and Abraxane^®^ in rhesus macaques and rats was confirmed by Student’s t-test indicating lack of significant difference in PK parameters. Accordingly, POXOL hl-PM is likely eligible to be tested in clinical trials through the 505(b)(2) regulatory pathway, which is an option for follow-on products of nanoformulated drugs if the newly formulated APIs have similar PK (bioequivalence) to the reference product [39]. In terms of the amount of data needed, the 505(b)(2) New Drug Application (NDA) falls somewhere in between the traditional 505(b)(1) NDA and the 505(j) abbreviated NDA (ANDA). The 505(b)(2) provides expediency by allowing reliance in part on the clinical data of safety and efficacy from the reference listed drug (RLD) and thereby shortens the time to approval and is more cost-effective.

The PET-CT biodistribution study in rhesus monkey showed a long blood circulation of POx, which can be attributed to the stealth properties of the shell-forming PMeOX block. Overall, our findings show that at the clinical dose of PTX, POXOL hl-PM is bioequivalent to Abraxane^®^ in rats and rhesus macaques. The hydrophilic PMeOx shell of POXOL hl-PM is biocompatible, making it a safer alternative to PEG-based systems, which have been shown to elicit an immune response in both animals and humans [40–42]. In summary, this work showcases comprehensive preclinical evaluations and highlights the potential of POXOL hl-PM to be developed into a clinical formulation.

## Acknowledgments

This work was supported by the National Cancer Institute (NCI) Alliance for Nanotechnology in Cancer (U54CA198999, Carolina Center of Cancer Nanotechnology Excellence) awarded to A.V.K. The studies at NCL were supported by federal funds from the National Cancer Institute, National Institutes of Health, under contract 75N91019D00024. We gratefully acknowledge kind assistance of Barry Neun, Jeff Clogston, and Marina A. Dobrovolskaia of the Nanotechnology Characterization Laboratory, US Frederick National Laboratory for Cancer Research, who assisted with the endotoxin assay and size distribution measurement and provided valuable insight. We would like to acknowledge the UNC Department of Comparative Medicine (DCM) Veterinarian Service team for animal procedures in rhesus macaque models, and the UNC Biomedical Research Imaging Center (BRIC) Imaging Facility for PET/CT imaging service. We also thank the UNC Lineberger Pathology Services Core for expert technical assistance with sciatic nerve staining for the histological assessment of neurotoxicity.

## Data availability

The raw/processed data required to reproduce these findings are available in the supplementary materials.

