## supplemental file for "Bioequivalence Assessment of High-Capacity Polymeric Micelle Nanoformulation of Paclitaxel and Abraxane^®^ in Rodent and Non-Human Primate Models Using a Stable Isotope Tracer Assay"

[#] Authors contributed equally on this study.

**Table S1.** Batch designation, drug loading, and POx block length of POXOL *hl*-PM formulations used in study.

| **Batch Designation** | **mg PTX/mg POx** | **Polymer block length** |
|---|---|---|
| Batch 1 (NCL 2014) | 10.0/12.5 | $P(MeOx_{33}$-b-$BuOx_{26}$-b-$MeOx_{45})$ |
| Batch 2 (NCL 2018) | 6.9/16.0 | $P(MeOx_{37}$-*b*-$BuOx_{21}$-*b*-$MeOx_{25})$ |
| Batch 3 (rat) | 6.9/16.0 | $P(MeOx_{37}$-*b*-$BuOx_{21}$-*b*-$MeOx_{25})$ |
| Batch 4 (rhesus macaque) | 120/300 | $P(MeOx_{37}$-*b*-$BuOx_{21}$-*b*-$MeOx_{25})$ |

**Table S2.** Physicochemical properties of POXOL *hl*-PM (Batch 2) (Loading Ratio 6.9/16.0 mg PTX/mg POx)

| Sample | **LE (%)** [a] | **LC (%)** [a] | **Drug conc. (g/L)** [a] | **Zeta Potential (mV)** [b] | | **DLS particle size data** | | | |
|---|---|---|---|---|---|---|---|---|---|
| | | | | **pH 7.0** | **pH 7.2** | **Z-avg, nm** [c] | **PDI** [c] | **Int-Peak, nm** [c] | **Vol-Peak, nm** [c] |
| Stock solution in saline | 92 | 26 | 5.88 | | | 30 ± 0 | 0.05 ± 0.01 | 32 ± 0 | 27 ± 0 |
| 10-fold dilution in saline | | | | | | 30 ± 0 | 0.07 ± 0.01 | 33 ± 0 | 26 ± 1 |
| 100-fold dilution in saline | | | | | | 32 ± 0 | 0.2 ± 0.02 | 34 ± 1 | 24 ± 1 |
| 10-fold dilution in water | | | | -3.0 ± 0.2 | -1.7 ± 0.4 | 32 ± 0 | 0.18 ± 0.03 | 34 ± 1 | 26 ± 1 |
| 100-fold dilution in water | | | | | | 98 ± 35 | 0.8 ± 0.11 | 677 ± 89 | 94 ± 1 |

[a] Determined by HPLC as described in Methods (section 2.2); [b] Determined by DLS as described in Methods (section 2.2) at NCL; [c] Determined by DLS as described in Methods (section 2.2) at NCL; The data present mean +/- standard deviation, z-average affective diameters along with the intensity and volume distribution diameters are presented.

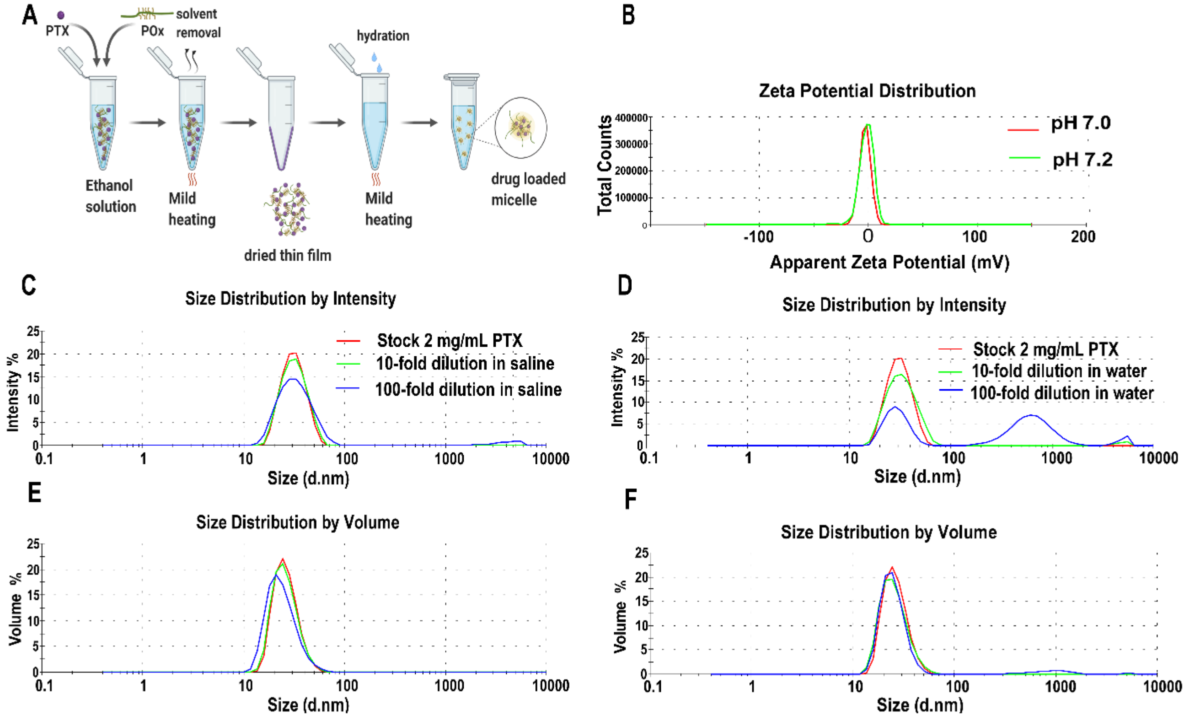


**Fig. S1 Preparation and physical characterization of POXOL *hl*-PM (Batch 2). (a)** Schematic of **POXOL *hl*-PM** preparation workflow (created with BioRender.com) **(b)** Zeta potential of **Batch 2** measured in water. Size distribution of **Batch 2** at different dilutions and dispersants measured by dynamic light scattering by **(c, d)** intensity and **(e, f)** volume (the details and quantitative data are presented in **Table S2**).

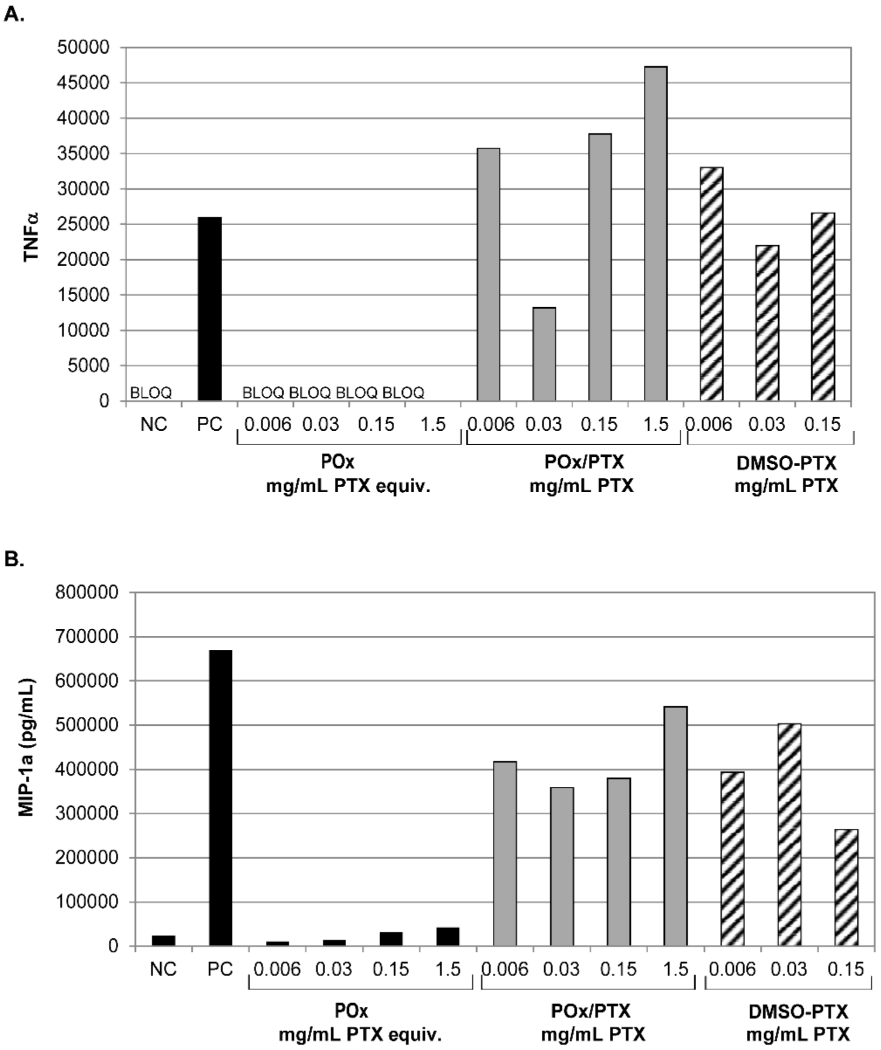


**Fig. S2 Cytokine induction by POXOL hl PM in RAW264.7 cells. (A)** Induction of TNF-α and **(B)** MIP-1a by POx polymer alone, POXOL hl PM (Batch 1) and unformulated paclitaxel in DMSO. PBS was used as the negative control (NC) and ultrapure K12 E. coli LPS was used as the positive control (PC). Data represent mean (n=2). POx/PTX indicates POXOL hl PM and DMSO-PTX indicates unformulated paclitaxel in DMSO. All concentrations shown in figure S2 are mg/mL PTX equivalent.

**Table S3:** Conversion factors used in the calculation of paclitaxel dose for PK study

| | **Clinical Dose# (mg/m$^2$)** | **Clinical Dose## (mg/kg)** | **mouse MTD* (mg/kg)** | **mouse MTD** (mg/m$^2$)** | **HED*** based on mouse MTD (mg/kg)** |
|---|---|---|---|---|---|
| **Abraxane®** | 260 | 7.0 | 90 | 270 | 7.3 |
| **Genexol®PM** | 435 | **11.8** | 50 | 150 | 4.1 |
| **POXOL hl-PM** | ND | ND | 150 | 450 | **12.2** |
| **Taxol** | 175 | 4.7 | 20 | 60 | 1.6 |

### MTD for given paclitaxel products previously determined in clinical trials [1, 2]; ##The conversion of human mg/m2 dose to mg/kg dose was performed by dividing the mg/m$^2$ dose by human $K_m$, 37 [3]; *Paclitaxel mouse MTD column determined previously with q4d x 4 regimen [4]; ** The conversion of mouse mg/kg dose to mg/m$^2$ dose was performed by multiplying the mg/m$^2$ dose by mouse $K_m$, 3[3]; *** mouse mg/kg dose was converted to human equivalent dose (HED), based on surface area, by dividing by 12.3 [3].

**Table S4:** Comparison of the total, protein-bound, encapsulated and free PTX PK parameters between POXOL hl PM (Batch 3) and Abraxane® for the study conducted using parallel study design in Sprague-Dawley rat model.

| | Animal code | $T_{max}$ | $C_{max}$ | $AUC_{all}$ | $AUC_{inf}$ | CL | $t_{1/2}$ | Lambda z | MTR | $V_{ss}$ |
|---|---|---|---|---|---|---|---|---|---|---|
| **PK Parameters** | | | | | | | | | | |
| | | h | ng/mL | ng*h/mL | ng*h/mL | mL/h*kg | h | $h^{-1}$ | H | mL/kg |
| **Total PTX** | | | | | | | | | | |
| **POXOL *hl*-PM** | **17** | 0.3 | 3019.8 | 5347.7 | 5530.8 | 2169.7 | 7.1 | 0.1 | 4.1 | 8824.5 |
| | **19** | 0.3 | 4155.2 | 7578.9 | 7719.6 | 1554.5 | 5.6 | 0.1 | 3.7 | 5749.7 |
| | **21** | 0.3 | 1417.4 | 3849.2 | 4228.4 | 2837.9 | 9.4 | 0.1 | 7.7 | 21772.1 |
| | **23** | 0.3 | 1865.7 | 4226.1 | 4322.1 | 2776.4 | 5.7 | 0.1 | 4.0 | 11175.0 |
| | **25** | 0.3 | 2535.4 | 4644.3 | 4731.8 | 2536.0 | 5.5 | 0.1 | 3.5 | 8943.9 |
| | **27** | 0.3 | 3330.3 | 7384.9 | 7482.1 | 1603.8 | 5.1 | 0.1 | 3.4 | 5494.5 |
| | **29** | 0.3 | 2934.9 | 5525.8 | 5675.9 | 2114.2 | 7.1 | 0.1 | 3.7 | 7857.0 |
| | **31** | 0.3 | 4068.1 | 6243.6 | 6350.9 | 1889.5 | 5.5 | 0.1 | 3.4 | 6428.7 |
| | **AVG** | 0.3 | 2915.9 | 5600.1 | 5755.2 | 2185.3 | 6.4 | 0.1 | 4.2 | 9530.7 |
| | **SE** | 0.0 | 341.7 | 490.1 | 475.9 | 175.6 | 0.5 | 0.0 | 0.5 | 1872.8 |
| **Abraxane®** | **1** | 0.3 | 2738.4 | 4641.9 | 4814.3 | 2492.6 | 7.4 | 0.1 | 4.4 | 10927.1 |
| | **3** | 0.3 | 1661.7 | 3840.7 | 3915.1 | 3065.1 | 5.2 | 0.1 | 4.1 | 12508.5 |
| | **5** | 0.3 | 1775.1 | 3473.8 | 3566.9 | 3364.2 | 5.8 | 0.1 | 4.4 | 14934.1 |
| | **7** | 0.3 | 2845.1 | 5311.2 | 5395.0 | 2224.3 | 5.1 | 0.1 | 3.5 | 7826.3 |
| | **9** | 0.3 | 1845.1 | 3666.5 | 3755.4 | 3195.4 | 5.8 | 0.1 | 4.1 | 13210.6 |
| | **11** | 0.3 | 2870.6 | 7007.6 | 7158.2 | 1676.4 | 5.4 | 0.1 | 4.3 | 7149.3 |
| | **13** | 0.3 | 3193.7 | 6110.0 | 6246.6 | 1921.0 | 5.8 | 0.1 | 4.0 | 7623.8 |
| | **AVG** | 0.3 | 2418.5 | 4864.5 | 4978.8 | 2562.7 | 5.8 | 0.1 | 4.1 | 10597.1 |
| | **SE** | 0.0 | 239.3 | 507.3 | 515.9 | 249.4 | 0.3 | 0.0 | 0.1 | 1173.2 |

| PK Parameters | | | | | | | | | | |
|---|---|---|---|---|---|---|---|---|---|---|
| | Animal code | $T_{max}$ h | $C_{max}$ ng/mL | $AUC_{all}$ ng*h/mL | $AUC_{inf}$ ng*h/mL | CL mL/h*kg | $t_{1/2}$ h | Lambda z $h^{-1}$ | MTR H | $V_{ss}$ mL/kg |
| **Protein bound PTX** | | | | | | | | | | |
| **POXOL *hl*-PM** | **17** | 0.3 | 2480.5 | 4310.8 | 4440.5 | 2702.4 | 6.6 | 0.1 | 3.9 | 10456.2 |
| | **19** | 0.3 | 3702.5 | 6466.7 | 6591.9 | 1820.4 | 5.8 | 0.1 | 3.6 | 6642.9 |
| | **21** | 0.3 | 1224.7 | 3475.7 | 3764.1 | 3188.0 | 9.0 | 0.1 | 7.0 | 22343.4 |
| | **23** | 0.3 | 1581.1 | 3449.8 | 3555.7 | 3374.9 | 6.2 | 0.1 | 4.4 | 14743.7 |
| | **25** | 0.3 | 2078.4 | 4116.4 | 4423.3 | 2712.9 | 10.1 | 0.1 | 6.0 | 16177.5 |
| | **27** | 0.3 | 2824.4 | 4809.6 | 4889.4 | 2454.3 | 5.2 | 0.1 | 3.6 | 8722.0 |
| | **29** | 0.3 | 2441.7 | 5646.2 | 5728.0 | 2095.0 | 5.9 | 0.1 | 3.2 | 6746.3 |
| | **31** | 0.3 | 3202.1 | 6024.8 | 6096.7 | 1968.3 | 5.0 | 0.1 | 3.1 | 6186.6 |
| | **AVG** | 0.3 | 2441.9 | 4787.5 | 4936.2 | 2539.5 | 6.7 | 0.1 | 4.3 | 11502.3 |
| | **SE** | 0.0 | 288.8 | 407.0 | 389.7 | 199.3 | 0.7 | 0.0 | 0.5 | 2040.7 |
| **Abraxane®** | **1** | 0.3 | 2136.4 | 3713.9 | 3817.1 | 3143.7 | 6.0 | 0.1 | 4.3 | 13464.2 |
| | **3** | 0.3 | 1228.2 | 3384.0 | 3440.2 | 3488.2 | 5.1 | 0.1 | 4.1 | 14199.4 |
| | **5** | 0.3 | 1469.6 | 2861.6 | 2929.0 | 4096.9 | 5.6 | 0.1 | 4.3 | 17589.9 |
| | **7** | 0.3 | 2410.9 | 4352.5 | 4412.0 | 2719.9 | 5.0 | 0.1 | 3.4 | 9252.9 |
| | **9** | 0.3 | 1581.9 | 3335.3 | 3406.9 | 3522.3 | 5.5 | 0.1 | 4.1 | 14318.4 |
| | **11** | 0.3 | 2380.1 | 6124.9 | 6410.9 | 1871.8 | 7.1 | 0.1 | 5.4 | 10185.5 |
| | **13** | 0.3 | 2315.4 | 4855.9 | 4969.4 | 2414.8 | 5.7 | 0.1 | 4.1 | 10020.7 |
| | **AVG** | 0.3 | 1931.8 | 4089.7 | 4197.9 | 3036.8 | 5.7 | 0.1 | 4.2 | 12718.7 |
| | **SE** | 0.0 | 185.8 | 423.0 | 449.7 | 285.3 | 0.3 | 0.0 | 0.2 | 1141.7 |

| | Animal code | $T_{max}$ h | $C_{max}$ ng/mL | $AUC_{all}$ ng*h/mL | $AUC_{inf}$ ng*h/mL | CL mL/h*kg | $t_{1/2}$ h | Lambda z $h^{-1}$ | MTR H | $V_{ss}$ mL/kg |
|---|---|---|---|---|---|---|---|---|---|---|
| **Encapsulated PTX** | | | | | | | | | | |
| **POXOL *hl*-PM** | **17** | 0.3 | 228.5 | 484.1 | 502.3 | 23888.1 | 4.1 | 0.2 | 4.5 | 106707.0 |
| | **19** | 0.5 | 123.4 | 487.1 | 495.3 | 24227.0 | 4.7 | 0.1 | 4.9 | 117720.5 |
| | **21** | 0.3 | 129.6 | 261.9 | 375.0 | 32003.7 | 18.7 | 0.0 | 17.9 | 573790.2 |
| | **23** | 0.5 | 187.5 | 435.9 | 406.8 | 29499.6 | 1.4 | 0.5 | 2.4 | 69769.4 |
| | **25** | 0.3 | 230.4 | 170.3 | 238.3 | 50353.1 | 1.7 | 0.4 | 1.2 | 60126.9 |
| | **27** | 2.0 | 668.8 | 1849.2 | 1857.9 | 6459.0 | 4.6 | 0.2 | 3.3 | 21609.9 |
| | **29** | 0.3 | 228.1 | -739.0 | -714.5 | -16794.5 | 4.2 | 0.2 | 1.4 | -23285.7 |
| | **31** | 0.3 | 545.1 | -552.0 | -539.3 | -22251.7 | 3.4 | 0.2 | 0.7 | -14925.1 |
| | **AVG** | 0.5 | 292.7 | 299.7 | 327.7 | 14785.2 | 5.5 | 0.2 | 4.5 | 114972.3 |
| | **SE** | 0.2 | 71.2 | 277.5 | 275.6 | 8841.9 | 2.0 | 0.1 | 2.0 | 68182.7 |
| **Abraxane®** | **1** | 0.3 | 349.5 | 342.7 | 358.0 | 33521.3 | 4.1 | 0.2 | 1.6 | 54534.4 |
| | **3** | 0.3 | 283.0 | 196.4 | 208.7 | 57504.9 | 5.0 | 0.1 | 1.9 | 107963.0 |
| | **5** | 0.3 | 119.7 | 302.4 | 320.5 | 37440.3 | 5.7 | 0.1 | 5.8 | 215288.8 |
| | **7** | 0.3 | 218.8 | 467.1 | 481.5 | 24922.8 | 4.5 | 0.2 | 4.0 | 100752.5 |
| | **9** | 0.5 | 180.2 | 38.1 | 42.4 | 283083.2 | 3.7 | 0.2 | 2.0 | 578285.6 |
| | **11** | 0.3 | 234.8 | 665.9 | 745.6 | 16095.2 | 1.6 | 0.4 | 1.1 | 18197.0 |
| | **13** | 0.3 | 640.7 | 1245.5 | 1270.9 | 9442.0 | 13.5 | 0.1 | 2.3 | 21570.3 |
| | **AVG** | 0.3 | 289.5 | 465.4 | 489.6 | 66001.4 | 5.5 | 0.2 | 2.7 | 156655.9 |
| | **SE** | 0.0 | 64.7 | 150.0 | 154.4 | 36659.4 | 1.4 | 0.0 | 0.6 | 74730.2 |

Table title: **PK Parameters**

| | Animal code | $T_{max}$ | $C_{max}$ | $AUC_{all}$ | $AUC_{inf}$ | CL | $t_{1/2}$ | Lambda z | MTR | $V_{ss}$ |
|---|---|---|---|---|---|---|---|---|---|---|
| | | h | ng/mL | ng*h/mL | ng*h/mL | mL/h*kg | h | $h^{-1}$ | H | mL/kg |
| **PK Parameters** | | | | | | | | | | |
| **Free PTX** | | | | | | | | | | |
| **POXOL *hl*-PM** | **17** | 0.3 | 310.8 | 499.2 | 510.1 | 23523.5 | 5.8 | 0.1 | 3.6 | 83645.4 |
| | **19** | 0.3 | 379.7 | 618.5 | 626.9 | 19142.8 | 5.3 | 0.1 | 3.2 | 60404.8 |
| | **21** | 0.5 | 122.1 | 315.7 | 335.7 | 35749.1 | 7.7 | 0.1 | 6.4 | 227561.0 |
| | **23** | 0.3 | 179.4 | 357.1 | 367.8 | 32626.7 | 6.2 | 0.1 | 4.4 | 143467.2 |
| | **25** | 0.3 | 226.6 | 450.8 | 482.9 | 24848.6 | 9.7 | 0.1 | 6.0 | 148098.2 |
| | **27** | 0.3 | 277.9 | 504.1 | 513.3 | 23376.2 | 5.3 | 0.1 | 3.6 | 84780.6 |
| | **29** | 0.3 | 265.1 | 543.1 | 552.3 | 21728.6 | 6.3 | 0.1 | 3.2 | 69809.5 |
| | **31** | 0.3 | 320.9 | 593.9 | 601.1 | 19964.4 | 5.0 | 0.1 | 3.2 | 64300.7 |
| | **AVG** | 0.3 | 260.3 | 485.3 | 498.8 | 25348.1 | 6.5 | 0.1 | 4.3 | 114060.3 |
| | **SE** | 0.0 | 29.2 | 37.7 | 36.4 | 2107.8 | 0.6 | 0.0 | 0.5 | 20652.5 |
| **Abraxane®** | **1** | 0.3 | 252.4 | 414.2 | 430.6 | 27868.5 | 7.1 | 0.1 | 4.6 | 126955.5 |
| | **3** | 0.3 | 150.5 | 352.3 | 356.4 | 33673.8 | 4.7 | 0.1 | 3.6 | 121391.9 |
| | **5** | 0.3 | 185.8 | 314.5 | 319.7 | 37533.0 | 5.2 | 0.1 | 3.7 | 137926.0 |
| | **7** | 0.3 | 215.5 | 413.4 | 418.1 | 28703.3 | 4.6 | 0.2 | 3.3 | 95344.1 |
| | **9** | 0.3 | 148.1 | 305.7 | 313.4 | 38288.7 | 5.9 | 0.1 | 4.1 | 157644.4 |
| | **11** | 0.3 | 255.7 | 650.6 | 681.3 | 17612.3 | 7.3 | 0.1 | 5.4 | 95956.2 |
| | **13** | 0.3 | 237.5 | 452.1 | 460.8 | 26039.3 | 5.5 | 0.1 | 3.8 | 97728.2 |
| | **AVG** | 0.3 | 206.5 | 414.7 | 425.8 | 29959.8 | 5.8 | 0.1 | 4.1 | 118992.3 |
| | **SE** | 0.0 | 17.3 | 44.4 | 47.6 | 2735.6 | 0.4 | 0.0 | 0.3 | 9081.2 |

**Table S5:** Comparison of the total, protein-bound, encapsulated and free PTX PK parameters between POXOL hl-PM (Batch 4) and Abraxane® for the study conducted using crossover study design in rhesus macaque.

| PK Parameters | | | | | | | | | | |
|---|---|---|---|---|---|---|---|---|---|---|
| | Animal code | $T_{max}$ | $C_{max}$ | $AUC_{all}$ | $AUC_{inf}$ | CL | t1/2 | Lambda z | MTR | $V_{ss}$ |
| | | h | ng/mL | ng*h/mL | ng*h/mL | mL/h*kg | h | $h^{-1}$ | H | mL/kg |
| **Total PTX** | | | | | | | | | | |
| **POXOL hl PM** | Bertha | 0.8 | 16587.0 | 20093.8 | 20235.5 | 593.0 | 8.2 | 0.1 | 3.8 | 2502.6 |
| | Gertrude | 0.5 | 13301.0 | 24302.7 | 24403.7 | 491.7 | 7.0 | 0.1 | 4.1 | 2119.2 |
| | **AVG** | 0.6 | 14944.0 | 22198.2 | 22319.6 | 542.4 | 7.6 | 0.1 | 4.0 | 2310.9 |
| | **SE** | 0.1 | 1643.0 | 2104.4 | 2084.1 | 50.6 | 0.6 | 0.0 | 0.1 | 191.7 |
| **Abraxane®** | Bertha | 0.5 | 14348.0 | 18984.1 | 19206.2 | 624.8 | 11.8 | 0.1 | 4.7 | 3339.4 |
| | Gertrude | 0.5 | 21451.0 | 25568.2 | 25674.2 | 467.4 | 6.7 | 0.1 | 3.6 | 1765.4 |
| | **AVG** | 0.5 | 17899.5 | 22276.2 | 22440.2 | 546.1 | 9.3 | 0.1 | 4.1 | 2552.4 |
| | **SE** | 0.0 | 3551.5 | 3292.0 | 3234.0 | 78.7 | 2.6 | 0.0 | 0.6 | 787.0 |
| **Protein-bound PTX** | | | | | | | | | | |
| **POXOL hl PM** | Bertha | 0.8 | 10261.4 | 13855.0 | 13898.7 | 863.4 | 6.9 | 0.1 | 3.7 | 3315.4 |
| | Gertrude | 0.5 | 8911.6 | 16290.0 | 16349.1 | 734.0 | 6.7 | 0.1 | 4.3 | 3257.8 |
| | **AVG** | 0.6 | 9586.5 | 15072.5 | 15123.9 | 798.7 | 6.8 | 0.1 | 4.0 | 3286.6 |
| | **SE** | 0.1 | 674.9 | 1217.5 | 1225.2 | 64.7 | 0.1 | 0.0 | 0.3 | 28.8 |
| **Abraxane®** | Bertha | 0.5 | 9531.1 | 13206.0 | 13287.3 | 903.1 | 7.1 | 0.1 | 4.6 | 4474.7 |
| | Gertrude | 0.5 | 14382.3 | 18397.6 | 18461.3 | 650.0 | 6.4 | 0.1 | 3.7 | 2497.4 |
| | **AVG** | 0.5 | 11956.7 | 15801.8 | 15874.3 | 776.6 | 6.8 | 0.1 | 4.1 | 3486.1 |
| | **SE** | 0.0 | 2425.6 | 2595.8 | 2587.0 | 126.6 | 0.4 | 0.0 | 0.5 | 988.7 |

| | Animal code | $T_{max}$ | $C_{max}$ | $AUC_{all}$ | $AUC_{inf}$ | CL | t1/2 | Lambda z | MTR | $V_{ss}$ |
|---|---|---|---|---|---|---|---|---|---|---|
| **PK Parameters** | | | | | | | | | | |
| | | h | ng/mL | ng*h/mL | ng*h/mL | mL/h*kg | h | $h^{-1}$ | H | mL/kg |
| **Encapsualted PTX** | | | | | | | | | | |
| **POXOL hl PM** | Bertha | 0.8 | 4718.0 | 4689.0 | 4817.6 | 2490.9 | 12.7 | 0.1 | 3.9 | 13833.5 |
| | Gertrude | 0.4 | 3885.0 | 4982.7 | 5019.8 | 2390.5 | 8.6 | 0.1 | 3.7 | 9948.1 |
| | **AVG** | 0.6 | 4301.5 | 4835.9 | 4918.7 | 2440.7 | 10.7 | 0.1 | 3.8 | 11890.8 |
| | **SE** | 0.2 | 416.5 | 146.8 | 101.1 | 50.2 | 2.1 | 0.0 | 0.1 | 1942.7 |
| **Abraxane®** | Bertha | 0.5 | 3259.0 | 3626.4 | 3674.1 | 3266.1 | 8.3 | 0.1 | 4.9 | 18302.4 |
| | Gertrude | 0.5 | 4995.0 | 4355.8 | 4390.5 | 2733.2 | 8.0 | 0.1 | 3.0 | 9490.5 |
| | **AVG** | 0.5 | 4127.0 | 3991.1 | 4032.3 | 2999.6 | 8.1 | 0.1 | 4.0 | 13896.4 |
| | **SE** | 0.0 | 868.0 | 364.7 | 358.2 | 266.5 | 0.1 | 0.0 | 0.9 | 4405.9 |
| **Free PTX** | | | | | | | | | | |
| **POXOL hl PM** | Bertha | 0.8 | 1607.6 | 2140.7 | 2146.5 | 5590.6 | 6.7 | 0.1 | 3.3 | 19375.9 |
| | Gertrude | 0.5 | 1690.4 | 2994.5 | 3002.9 | 3996.2 | 6.5 | 0.1 | 3.7 | 15504.3 |
| | **AVG** | 0.6 | 1649.0 | 2567.6 | 2574.7 | 4793.4 | 6.6 | 0.1 | 3.5 | 17440.1 |
| | **SE** | 0.1 | 41.4 | 426.9 | 428.2 | 797.2 | 0.1 | 0.0 | 0.2 | 1935.8 |
| **Abraxane®** | Bertha | 0.5 | 1557.9 | 2130.9 | 2148.5 | 5585.2 | 11.1 | 0.1 | 4.3 | 26819.3 |
| | Gertrude | 0.5 | 2073.7 | 2790.1 | 2800.6 | 4284.9 | 6.6 | 0.1 | 3.7 | 16522.4 |
| | **AVG** | 0.5 | 1815.8 | 2460.5 | 2474.5 | 4935.0 | 8.8 | 0.1 | 4.0 | 21670.9 |
| | **SE** | 0.0 | 257.9 | 329.6 | 326.0 | 650.2 | 2.3 | 0.0 | 0.3 | 5148.5 |

**Table S6: Freeze-thaw control study of POXOL hl PM and Abraxane®.** Presented are the analytical data for the freeze-thaw control study for POXOL *hl*-PM and Abraxane®

| Controls | Filtrate PTX (ng/mL) | Reservoir PTX (ng/mL) | %Unbound PTX[1] | %Bound[2] | Filtrate PTX_C13 (ng/mL) | Reservoir PTX_C13 (ng/mL) | %Unbound PTX_C13[3] | %Bound PTX_C13[4] | Unencapsulated[5] | Encapsulated[6] | %Release[7] |
|---|---|---|---|---|---|---|---|---|---|---|---|
| POXOL fresh | 579 | 5731 | 10.1 | 90 | 11.2 | 96 | 11.7 | 88 | 4958 | 773 | 87 |
| POXOL fresh | 623 | 5518 | 11.3 | 89 | 12 | 105 | 11.4 | 89 | 5468 | 50 | 99 |
| POXOL fresh | 586 | 5776 | 10.1 | 90 | 12.5 | 102 | 12.2 | 88 | 4803 | 973 | 83 |
| Average (STD) | | | | | | | | | | | 90 (7) |
| POXOL frozen | 533 | 5484 | 9.7 | 90 | 12.9 | 108 | 11.9 | 88 | 4474 | 1009 | 82 |
| POXOL frozen | 515 | 5471 | 9.4 | 91 | 12.6 | 107 | 11.8 | 88 | 4381 | 1090 | 80 |
| POXOL frozen | 524 | 5921 | 8.9 | 91 | 12.4 | 109 | 11.4 | 89 | 4600 | 1320 | 78 |
| Average (STD) | | | | | | | | | | | *80 (2) |
| Abraxane fresh | 500 | 5539 | 9 | 91 | 11.7 | 100 | 11.7 | 88 | 4272 | 1268 | 77 |
| Abraxane fresh | 639 | 5570 | 11.5 | 89 | 11.6 | 111 | 10.5 | 90 | 6092 | -522 | 109 |
| Abraxane fresh | 557 | 5434 | 10.2 | 90 | 11.7 | 102 | 11.5 | 89 | 4857 | 577 | 89 |
| Average (STD) | | | | | | | | | | | 92 (13) |
| Abraxane frozen | 497 | 5208 | 9.5 | 90 | 12.7 | 113 | 11.3 | 89 | 4397 | 811 | 84 |
| Abraxane frozen | 455 | 4757 | 9.6 | 90 | 13.1 | 112 | 11.7 | 88 | 3884 | 873 | 82 |
| Abraxane frozen | 451 | 4512 | 10 | 90 | 14 | 113 | 12.4 | 88 | 3629 | 883 | 80 |
| Average (STD) | | | | | | | | | | | *82 (2) |

[1] = Filtrate PTX / Reservoir PTX × 100, [2] = 100 – %Unbound PTX, [3] = (Filtrate PTX_C13 / Reservoir PTX_C13) × 100, [4] = 100 – %Unbound PTX_C13, [5] = Filtrate PTX / (1 – (%Bound PTX_C13) / 100)), [5] = Reservoir PTX – Unencapsulated, [6] = (Unencapsulated / Reservoir PTX) × 100,
*Not significantly different by Student's t-test

**Table S7: Complete blood count.** Results of the complete blood count performed on rhesus macaques following treatment with the test (POXOL hl PM) and reference (Abraxane) formulations.

| **Cell Count Normal range**[1] | **Rhesus macaque 1** | | | | | **Rhesus macaque 2** | | | | |
|---|---|---|---|---|---|---|---|---|---|---|
| | Baseline | POXOL hl PM (1 week) | % change POXOL hl PM | Abraxane (1 week) | % change Abraxane | Baseline | Abraxane (1 week) | % change Abraxane | POXOL hl PM (1 week) | % change POXOL hl PM |
| **WBC ($10^9$/L) 5.7 – 21** | 5.79 | 11.19 | 93.3 | 16.19 | 180 | 5.98 | 14.82 | 148 | 2.27 | -62 |
| **LYM ($10^9$/L) 4.3 – 9.24** | 4.73 | 6.23 | 31.7 | 9.35 | 97.7 | 0.1 | 3.41 | 3310 | 2.28 | 2180 |
| **MON ($10^9$/L) 0.04 – 0.72** | 0.33 | 1.42 | 330.3 | 2.32 | 603 | 4.05 | 0.08 | -98.1 | 0.02 | -99.5 |
| **GRA ($10^9$/L) 1.9 – 6.24** | 0.73 | 3.55 | 386.3 | 4.52 | 519.2 | 1.83 | 11.32 | 518.6 | 0.07 | -96.2 |
| **RBC ($10^{12}$/L) 4.8 – 6.3** | 5.2 | 5.07 | -2.5 | 4.53 | -13 | 5.44 | 5.24 | -3.7 | 4.75 | -12.7 |
| **HGB (g/dL) 8 - 15** | 11.9 | 11.5 | -3.4 | 10.2 | -14.3 | 11.3 | 11.9 | 5.3 | 10 | -11.5 |
| **HCT (%) 30 - 44** | 36.55 | 36.12 | -1.2 | 31.77 | -13.1 | 36.39 | 35.62 | -2.1 | 30.69 | -15.7 |
| **MCV (fL) 50 - 90** | 70 | 71 | 1.4 | 70 | 0 | 67 | 68 | 1.5 | 65 | -3 |
| **MCH (pg) 12 - 13** | 22.9 | 22.6 | -1.3 | 22.5 | -1.8 | 20.8 | 22.8 | 9.6 | 21 | 1 |
| **MCHC (g/dL) 30 - 36** | 32.6 | 31.8 | -2.5 | 32 | -1.8 | 31.1 | 33.5 | 7.7 | 32.5 | 4.5 |
| **PLT ($10^9$/L) 200 - 600** | 167 | 178 | 6.6 | 73 | -56.3 | 111 | 349 | 214.4 | 227 | 104.5 |

**[1]Abbreviation of types of blood cells and blood values in table S9: white blood cell (WBC), lymphocyte (LYM), monocyte (MON), granulocyte (GRA), red blood cell (RBC), hemoglobin (HGB), hematocrit (HCT), mean corpuscular volume (MCV), mean corpuscular hemoglobin (MCH), mean corpuscular hemoglobin concentration (MCHC), and platelet (PLT)**

**Table S8: Blood chemistry panel.** Results of the blood chemistry panel performed on rhesus macaques following treatment with the test (POXOL hl-PM) and reference (Abraxane®) formulations.

| Blood chemistry Normal range[1] | Rhesus macaque 1 | | | | | Rhesus macaque 2 | | | | |
|---|---|---|---|---|---|---|---|---|---|---|
| | Baseline | POXOL hl PM (1 week) | % change POXOL hl PM | Abraxane (1 week) | % change Abraxane | Baseline | Abraxane (1 week) | % change Abraxane | POXOL hl PM (1 week) | % change POXOL |
| **ALB (g/dL) 2.4 – 3.3** | 4.3 | 3.8 | -11.6 | 3.6 | -16.3 | 4.6 | 3.6 | -21.7 | 4.2 | -8.7 |
| **ALP (U/L) 100 - 250** | 62 | 115 | 85.5 | 79 | 27.5 | 86 | 138 | 60.5 | 83 | -3.5 |
| **ALT (U/L) 10 - 100** | 24 | 34 | 41.7 | 24 | 0 | 59 | 30 | -49.2 | 22 | -62.7 |
| **AMY (U/L) 1000 - 2500** | 289 | 218 | -25.6 | 312 | 8 | 306 | 305 | -0.3 | 298 | -2.6 |
| **TBIL (mg/dL) 0 - 1** | 0.3 | 0.3 | 0 | 0.4 | 33.3 | 0.3 | 0.3 | 0 | 0.4 | 33.3 |
| **BUN (mg/dL) 10 - 22** | 14 | 15 | 7.1 | 13 | -7.1 | 19 | 24 | 26.3 | 13 | -31.6 |
| **Ca2+ (mg/dL) 7.2 - 11.5** | 9.2 | 9.6 | 4.3 | 9.2 | 0 | 9.2 | 9.9 | 7.6 | 9.7 | 5.4 |
| **PHOS(mg/dL) 3 - 5.5** | 4.6 | 3.4 | -26 | 4.7 | 2.2 | 3.2 | 4.5 | 40.6 | 2.6 | -18.8 |
| **CRE (mg/dL) 0.1 - 2** | 1 | 1 | 0 | 0.8 | -20 | 1 | 0.9 | -10 | 0.9 | -10 |
| **GLU(mg/dL) 70 - 120** | 130 | 93 | -28.5 | 114 | -12.3 | 121 | 181 | 49.6 | 131 | 8.3 |
| **Na+ (mmo/L) 138 - 150** | 137 | 132 | -3.7 | 138 | 0.7 | 138 | 131 | -5.1 | 134 | -2.9 |
| **K+ (mmo/L) 3.5 – 5.5** | 3.7 | 4.3 | 16.2 | 4.1 | 10.8 | 3.8 | 5.1 | 34.2 | 3.6 | -5.3 |
| **TP (g/dL) 5 - 7** | 6.5 | 6.7 | 3.1 | 5.9 | -9.2 | 7 | 6.8 | -2.9 | 6.7 | -4.3 |
| **GLOB (g/dL) 2 - 4** | 2.2 | 2.9 | 0.3 | 2.3 | 4.5 | 24 | 3.3 | 37.5 | 2.5 | 4.2 |

**[1]Abbreviation of blood chemistry parameters in table S8: albumin (ALB), alkaline phosphatase (ALP), alanine aminotransferase (ALT), amylase (AMY), total bilirubin (TBIL), blood urea nitrogen (BUN), calcium (Ca+), phosphate (PHOS), creatinine (CRE), glucose (GLU), sodium (Na+), potassium (K+), total protein (TP), and globulin (GLOB)**
